# *Trypanosoma brucei* PolIE suppresses telomere recombination

**DOI:** 10.1101/2021.05.07.443201

**Authors:** Maiko Tonini, M. A. G. Rabbani, Marjia Afrin, Bibo Li

## Abstract

Telomeres are essential for genome integrity and stability. In *T. brucei* that causes human African trypanosomiasis, the telomere structure and telomere proteins also influence the virulence of the parasite, as its major surface antigen involved in the host immune evasion is expressed exclusively from loci immediately upstream of the telomere repeats. However, telomere maintenance mechanisms are still unclear except that telomerase-mediated telomere synthesis is a major player. We now identify PolIE as an intrinsic telomere complex component. We find that depletion of PolIE leads to an increased amount of telomere/subtelomere DNA damage, an elevated rate of antigenic variation, and an increased amount of telomere T-circles and C-circles, indicating that PolIE suppresses telomere recombination and helps maintain telomere integrity. In addition, we observe much longer telomere G-rich 3’ overhangs in PolIE-depleted cells, which is not dependent on telomerase. Furthermore, the level of telomere DNA synthesis is slightly increased in PolIE-depleted cells, which is dependent on telomerase. Therefore, we identify PolIE as a major player for telomere maintenance in *T. brucei*.

## Introduction

Telomeres are nucleoprotein complexes at chromosome ends and are essential for genome integrity and chromosome stability (1, 2). In most eukaryotic organisms, telomeres consist of simple repetitive sequences with the G-rich strand going 5’ to 3’ towards the chromosome ends (3). Telomere length maintenance involves several steps, and it is important to coordinate between the telomere G-strand extension and C-strand fill-in. In the S phase, the chromosome internal portion of the telomere is replicated by conventional DNA polymerases, which are incapable of fully replicating the ends of linear DNA molecules, resulting in a so-called “end replication problem” (4). In most eukaryotic organisms, a specialized reverse transcriptase, telomerase, synthesizes the G-rich strand telomere DNA *de novo* (5, 6). Telomerase contains a protein subunit, TERT, which contains the catalytic activity, and an RNA subunit, TR, which provides a short sequence template for telomere synthesis (5, 6). Telomerase uses a G-rich single-stranded 3’ DNA end as its substrate (7). Hence, the telomere single-stranded 3’ G-rich overhang structure is important for telomerase-mediated telomere synthesis (8–11). The telomere G-overhang is also critical for the formation of the telomere T-loop structure (12, 13), which in turn helps protect the telomere from illegitimate nucleolytic degradation and DNA damage repair processes (2). Since leading strand DNA synthesis results in blunt-ended DNA products, exonucleases are required to resect these telomere 5’ ends to generate the telomere 3’ G-overhang structure (14). In human cells, the Apollo exonuclease is recruited to the telomere by the duplex telomere DNA binding factor, TRF2 (15, 16), to blunt-ended telomere ends to resect the telomere 5’ ends (17). Subsequently, EXO1 takes over the 5’ to 3’ end degradation process to generate longer telomere G-overhangs (17). After telomerase-mediated telomere G-strand synthesis, the telomere C-strand is filled-in by the DNA Primase-Polymerase alpha (18, 19). In addition, the coordination between the telomere G-strand synthesis and C-strand fill-in is mainly regulated by proteins binding the single-stranded telomere G-overhang, including POT1 (20, 21) and the CST complex (CTC1/STN1/TEN1 in mammals and CDC13/STN1/TEN1 in budding yeast) (22–26). Therefore, the telomere G-overhang can be elongated by telomerase (7) and by exonucleases that degrade the telomere 5’ end sequences (17). On the other hand, it can be shortened when the telomere C-strand is filled-in by the DNA Primase-Polymerase alpha (27).

*Trypanosoma brucei* is a protozoan parasite that causes human African trypanosomiasis. While proliferating in its mammalian host, *T. brucei* sequentially expresses immunologically distinct VSGs, its major surface antigen, to evade the host immune response. *T. brucei* has a large *VSG* gene pool and all *VSGs* are located at subtelomeric regions (28). However, only those in VSG expression sites (ESs) can be expressed in a strictly monoallelic manner (29). ESs are polycistronic transcription units, and *VSG* is the last gene in each ES, located immediately upstream of the telomere repeats (30). VSG switching is frequently mediated by DNA recombination and sometimes through transcriptional switches (31, 32).

Telomerase-mediated telomere synthesis is the predominant mechanism of telomere maintenance in *T. brucei* (33–35). *T. brucei* telomeres also form a T-loop structure (13). In addition, *T. brucei* telomere has a short 3’ single-stranded G-rich overhang (10, 11). The telomere structure and telomere proteins are not only essential for *T. brucei* genome stability and cell proliferation (36) but are also essential for monoallelic *VSG* expression (37–40) and influence the VSG switching frequency (41–44). However, detailed *T. brucei* telomere synthesis mechanisms are still not clear. We previously found that *T. brucei* has very short telomere G-overhangs (∼12 nts) (10) and the telomerase activity is essential for the telomere G-overhang structure (11), suggesting that resection of the telomere 5’ end by an exonuclease, which has not been identified in *T. brucei* yet, is minimum and/or the telomere C-strand fill-in step is efficient. So far, no protein has been identified to bind specifically to the single-stranded telomere DNA in *T. brucei*, and how telomere C-strand fill-in is achieved is unknown.

In this study, we performed both Proteomics of Isolated Chromatin Segments (PICh) of the telomere chromatin and affinity pull-down of the telomere protein complex. We have identified PolIE as an intrinsic component of the *T. brucei* telomere complex. We show that PolIE is essential for cell proliferation and helps maintain genome integrity. Importantly, PolIE plays critical roles in maintaining telomere stability. First, in cells depleted of PolIE, we detected a much higher amount of telomere T-circles and C-circles, indicating that PolIE normally suppresses telomere homologous recombination (HR). Second, PolIE-depleted cells have much longer telomere G-overhangs, which is not telomerase-dependent. Third, a higher level of telomerase-dependent telomere synthesis is detected in PolIE-depleted cells, indicating that PolIE normally inhibits telomerase-mediated telomere G-strand extension.

## Results

### Identify T. brucei telomere components by PICh and TbTRF/TbTIF2 IP

We previously identified *Tb*TRF as a duplex telomere DNA binding factor (36) and *Tb*TIF2 as a *Tb*TRF interacting factor (42). We have also shown that telomere proteins play important roles in VSG regulation in *T. brucei* (37–43, 45, 46). To identify additional telomere proteins, we first pulled down the known telomere protein complex. We tagged both *Tb*TRF and *Tb*TIF2 with FLAG-HA-HA (F2H) at one of their respective endogenous loci and replaced the remaining allele with selectable markers to generate the procyclic form (PF, the insect stage) *T. brucei TbTRF*^F2H+/-^ *TbTIF2*^+F2H/-^ strain. The nuclear extracts prepared from this strain and WT cells (*TbTRF*^+/+^ *TbTIF2*^+/+^, as a negative control) were first immunoprecipitated (IPed) with the FLAG monoclonal antibody M2 (Sigma) then with the HA monoclonal antibody 12CA5 (MSKCC Antibody & Bioresource Core Facility), and the final IP products were analyzed by mass spectrometry (Cleveland Clinic Lerner Research Institute Proteomics and Metabolomics Core) (Fig. S1, left). Proteins identified from WT cells were considered as the background noise and were eliminated from those identified in the *TbTRF*^F2H+/-^ *TbTIF2*^+F2H/-^ cells. To identify all proteins associated with the *T. brucei* telomere chromatin in an unbiased manner, we also performed Proteomics of Isolated Chromatin segments (PICh) (47). Using an LNA probe containing either the (TTAGGG)^4^ sequence or a scrambled sequence (as a negative control), we pulled down the telomere chromatin and analyzed all its protein components by mass spectrometry (Fig. S1, right). Proteins identified using the control probe were considered as the background noise and removed from those identified using the (TTAGGG)^4^ LNA probe.

Both IP and PICh identified several hundred proteins, and 283 proteins were identified in both approaches (Fig. 1A). Among these, DNA polymerase IE (Tb927.11.5550) (48, 49) is one of the most abundant proteins. In addition, we have identified *Tb*TRF (36), *Tb*TIF2 (42), *Tb*RAP1 (37), NUP98, NUP96, NUP152, SMC1, SMC3, SMC4, PPL2 (48), and TelAP1 (48), all core histones, H3v, and H2Bv, etc. Most of these proteins have been identified to associate with the telomere chromatin previously, confirming that our approaches are successful.

**Figure 1.**
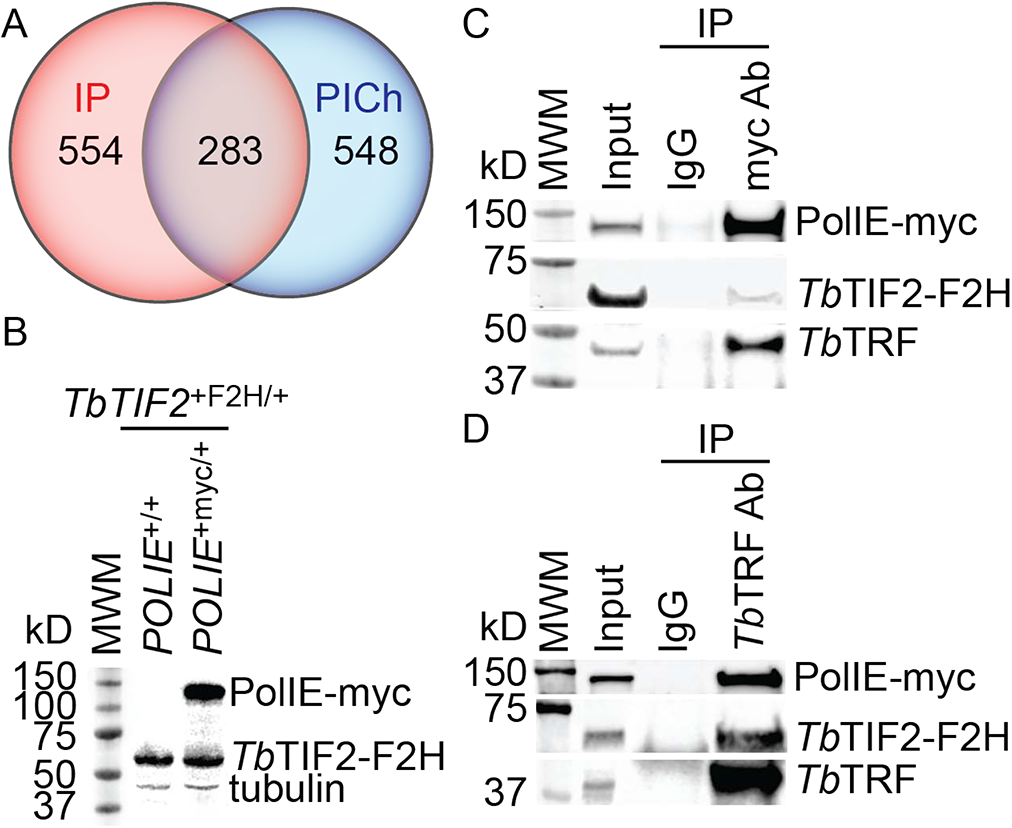
PolIE interacts with *Tb*TRF and *Tb*TIF2. (A) Venn diagram showing the number of protein candidates identified in IP of the *Tb*TRF/*Tb*TIF2 protein complex and in PICh. (B) Western blotting showing the expression of *Tb*TIF-F2H and PolIE-myc in the indicated strains. (C) and (D) IP using the myc Ab 9E10 (MSKCC Antibody & Bioresource Core Facility) (C) and a *Tb*TRF rabbit Ab (36) (D) and IgG (as a negative control) in *TbTIF2*^+F2H/+^ *POLIE*^+myc/+^ cells. Western analyses were performed using the myc Ab 9E10, the HA antibody (HA probe, Santa Cruz Biotechnologies), and a *Tb*TRF chicken Ab (37).

### T. brucei DNA Polymerase IE associates with the telomere chromatin

Although PolIE was previously identified in a pull-down using a telomere oligo and in a *Tb*TRF IP (48), it has not been directly confirmed that PolIE is a telomere protein (48, 49). To verify that PolIE is indeed an intrinsic component of the telomere complex, we first C-terminally tagged one *POLIE* endogenous allele with 13x myc in the bloodstream form (BF, the mammalian infectious stage) *T. brucei* cells where one *TbTIF2* endogenous allele is C-terminally tagged with an F2H epitope (Fig. 1B). *POLIE*^+/+^, *POLIE*^+myc/+^, *POLIE*^+/-^, and *POLIE*^+myc/-^ cells grew with nearly identical rate (Fig. S2A), indicating that PolIE-myc is functional. In PolIE-myc and *Tb*TIF2-F2H expressing cells, IP of PolIE-myc using the myc monoclonal antibody 9E10 (MSKCC Antibody & Bioresource Core Facility) pulled down *Tb*TIF2-F2H and *Tb*TRF in addition to PolIE-myc (Fig. 1C). Similarly, PolIE-myc and *Tb*TIF2-F2H were present in the *Tb*TRF IP product when a *Tb*TRF rabbit antibody (36) was used (Fig. 1D). Therefore, PolIE interacts with *Tb*TRF and *Tb*TIF2 in *T. brucei* cells.

Subsequently, we performed ChIP experiment using the myc antibody 9E10 and observed that PolIE-myc, similar to *Tb*TRF, is associated with the telomere chromatin but not the tubulin chromatin (Fig. 2, A, B; (36) Fig. S2B). Furthermore, we performed Immunofluorescent (IF) analysis in PolIE-myc expressing cells using the myc antibody and *Tb*TRF antibody (36), where *Tb*TRF is used as a telomere marker (36). We observed that PolIE-myc is colocalized with *Tb*TRF throughout the cell cycle (Fig. 2C). Therefore, PolIE is an intrinsic component of the telomere complex.

**Figure 2.**
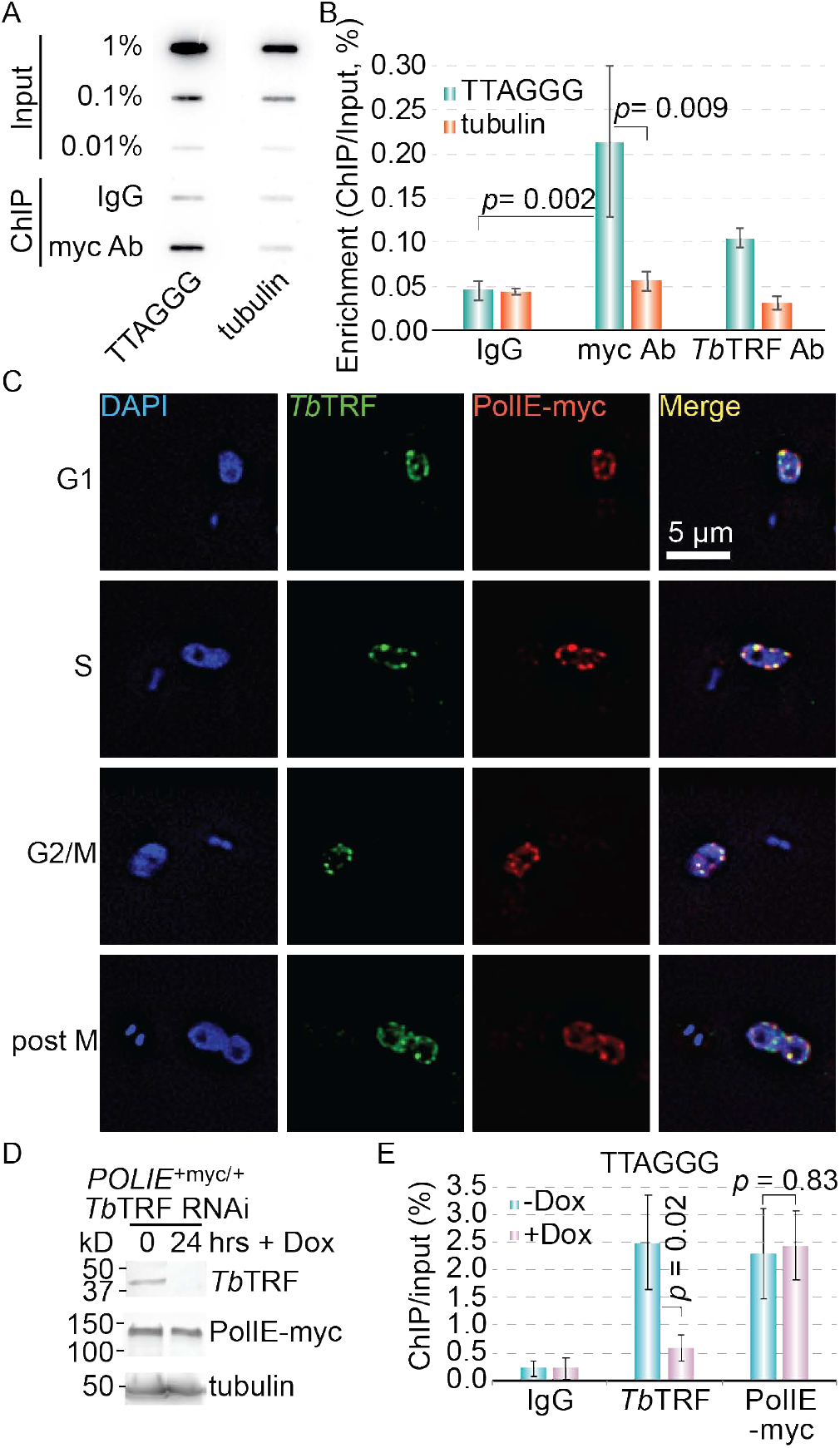
PolIE is enriched at the telomere. (A) ChIP using the myc Ab 9E10, a *Tb*TRF rabbit antibody (36), and IgG (as a negative control) in *POLIE*^+myc/+^ cells. The ChIP products were detected in Southern slot blots using a telomere and a tubulin probe. (B) Quantification results of PolIE-myc and *Tb*TRF ChIP in *POLIE*^+myc/+^ cells. Average enrichment (ChIP/Input) was calculated from three or four independent experiments. (C) IF in *POLIE*^+myc/+^ cells using the myc Ab 9E10 and a *Tb*TRF rabbit Ab. DNA was stained with DAPI. All panels are of the same scale and the size bar is shown in one panel. Representative cells in various cell cycle stages are shown. (D) Western blotting showing depletion of *Tb*TRF in *POLIE*^+myc/+^ *Tb*TRF RNAi cells after a 24-hr induction. A *Tb*TRF rabbit Ab, the myc Ab 9E10, and the tubulin Ab TAT-1 (81) were used. (E) Quantification results of PolIE-myc and *Tb*TRF ChIP in *POLIE*^+myc/+^ *Tb*TRF RNAi cells before (-Dox) and after (+Dox) the induction of *Tb*TRF RNAi. Average enrichment (ChIP/Input) was calculated from three independent experiments. In this and other figures, error bars represent standard variation. *P* values of unpaired *t*-tests are shown in (B) and (E).

We further investigated whether the association of PolIE with the telomere chromatin depends on *Tb*TRF, which has a duplex telomere DNA binding activity (36). ChIP was performed in PolIE-myc expressing *Tb*TRF RNAi cells before and after induction of *Tb*TRF RNAi for 24 hrs using the myc antibody 9E10, a *Tb*TRF rabbit antibody (as a positive control) (36), or IgG (as a negative control). ChIP products were then hybridized with a telomere and a tubulin probe (as a negative control) (Fig. 2E; Fig. S2, C, D). *Tb*TRF was successfully depleted and the association of *Tb*TRF and the telomere chromatin was abolished upon induction of *Tb*TRF RNAi (Fig. 2, D, E; Fig. S2C). However, PolIE still remained at the telomere after depletion of *Tb*TRF (Fig. 2E; Fig. S2, C, D). Therefore, although PolIE and *Tb*TRF interact, PolIE is localized to the telomere independent of *Tb*TRF.

### PolIE is essential for cell proliferation and important for cells to cope with DNA damage assaults

PolIE is an A type DNA polymerase that has a DNA polymerase domain very similar to that in mammalian Polθ (49). However, Polθ has both a C-terminal DNA polymerase domain and an N-terminal helicase domain (50), the latter of which is missing in PolIE (49). Biochemical studies have shown that Polθ has a weak translesion DNA synthesis activity, while Polθ homologues have been shown to be an important factor of Microhomology-Mediated End Joining (MMEJ) that can help repair DNA double strand breaks (DSBs) (51, 52). To examine potential functions of PolIE in DNA damage repair, we first established inducible PolIE RNAi strains that also express PolIE-myc from its endogenous locus. Significant depletion of PolIE-myc and growth arrest by 24 hrs (Fig. 3, A, B) were observed upon induction of PolIE RNAi, confirming that PolIE is essential for cell proliferation (49).

**Figure 3.**
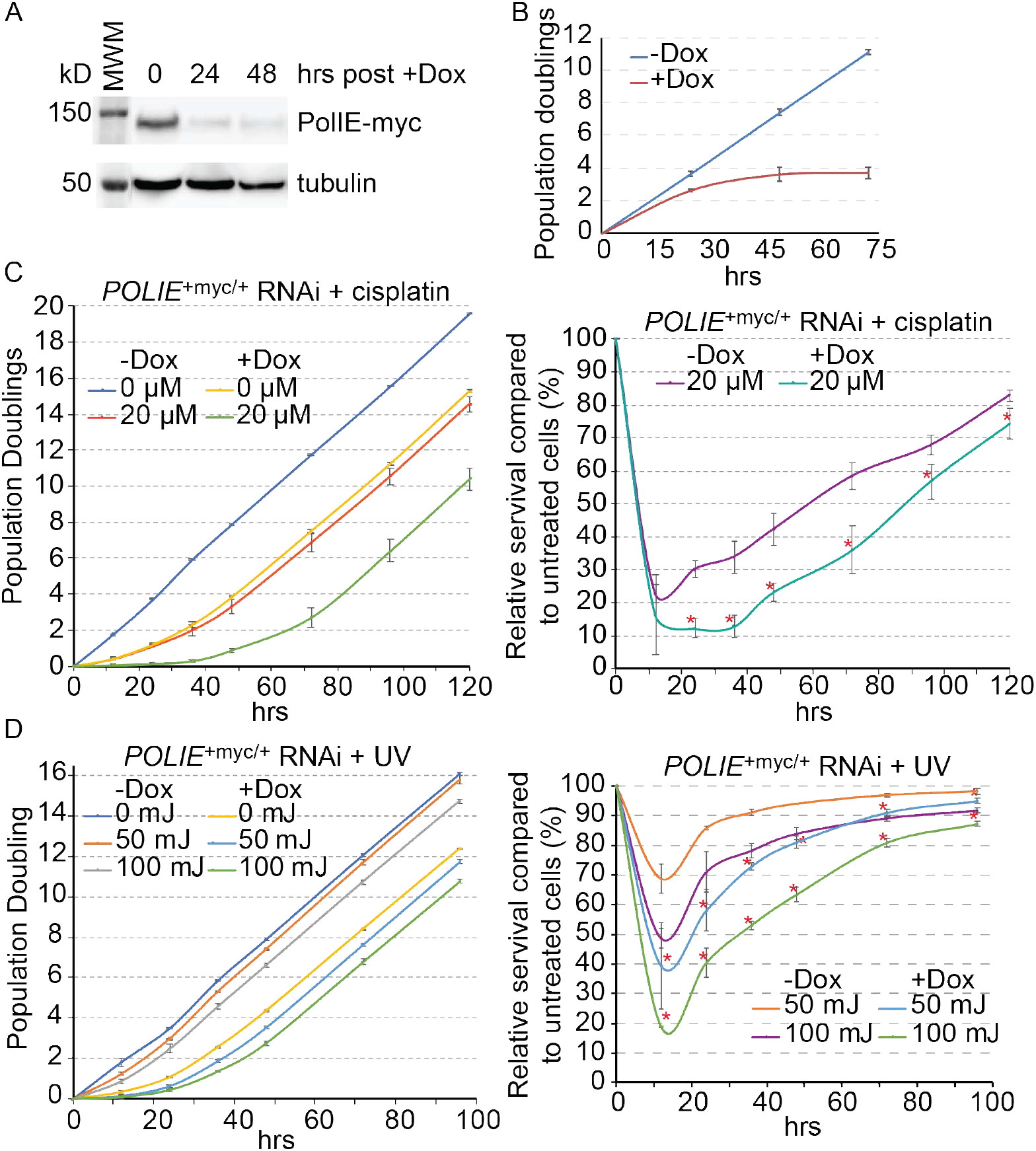
PolIE-depleted cells are hypersensitive to cisplatin treatment and UV irradiation. (A) Western blotting showing depletion of PolIE-myc in *POLIE*^+myc/+^ RNAi cells. The myc Ab 9E10 and the tubulin Ab TAT-1 (81) were used. (B) Growth curves of *POLIE*^+myc/+^ RNAi cells before (-Dox) and after (+Dox) the induction of RNAi. (C) *POLIE*^+myc/+^ RNAi cells incubated with or without doxycycline for 12 hrs were treated with and without 20 µM cisplatin for 1 hr before cells were washed free of doxycycline and cisplatin. Subsequently, cell growth was monitored (left) and relative growth (treated/untreated) was calculated from three independent experiments and shown on the right. (D) *POLIE*^+myc/+^ RNAi cells incubated with or without doxycycline for 12 hrs were irradiated with and without 50 mJ and 100 mJ UV before cells were washed free of doxycycline. Subsequently, cell growth was monitored (left) and relative growth (treated/untreated) was calculated from three independent experiments and shown on the right. Asterisks indicate significant difference between relative growth of uninduced and induced cells.

Mammalian cells lacking Polθ are hypersensitive to DNA damaging agents that cause DSBs (50). To examine whether PolIE has a similar function in *T. brucei*, we first examined whether PolIE-depleted cells were more sensitive to EMS. BF PolIE RNAi cells were induced for 0 and 24 hrs followed by treatment with 2 mM EMS (Sigma) for 2 hrs. Subsequently, cells were washed free of doxycycline and EMS and plated in 96-well dishes. We first calculated the plating efficiency (number of survival clones/total number of wells). Relative survival rates (plating efficiency of EMS treated/plating efficiency of untreated) were then calculated for induced and un-induced PolIE RNAi cells. As shown in Fig. S2E, cells depleted of PolIE exhibited a significantly reduced survival rate than un-induced cells. Therefore, PolIE-depleted cells are more sensitive to DNA damage reagent EMS, which is consistent with the previous observation that PolIE-depleted cells were slightly more sensitive to MMS treatment (49)

We further investigated whether depletion of PolIE resulted in hyper sensitivity to UV and cisplatin (causes interstrand crosslink, ICL) in *T. brucei*. PolIE RNAi cells were induced by doxycycline for 24 hrs and incubated with or without 20 µM cisplatin for 1 hr or irradiated with 0, 50 mJ, and 100 mJ of UV light. Subsequently, cells were washed extensively to remove doxycycline and cisplatin and continuously cultured, and cell growth was monitored. As shown in Fig. 3C, cells treated with cisplatin had a poorer relative growth (treated/untreated) in PolIE-depleted cells than in uninduced cells (Fig. 3C, right). Similarly, cells irradiated with UV exhibited much poorer growth than un-irradiated cells (Fig. 3D, left), and the relative growth (irradiated/un-irradiated) is significantly poorer for PolIE-depleted cells compared to un-induced cells (Fig. 3D, right). Therefore, depletion of PolIE results in hypersensitivity to both UV irradiation and cisplatin treatment in *T. brucei*.

### Loss of PolIE results in an increased amount of DNA damage at the telomere vicinity and an elevated VSG switching frequency

The above-described observations suggest that PolIE may be important for maintaining telomere integrity. Therefore, we examined whether depletion of PolIE caused an increased amount of telomere DNA damage. Western analysis showed that depletion of PolIE by RNAi did not affect the protein levels of *Tb*TRF or *Tb*RAP1 (Fig. 4A). However, we observed an increased amount of γH2A, a marker of damaged DNA (53), upon depletion of PolIE (Fig. 4A). IF using a γH2A rabbit antibody (43) showed that more than 90% of PolIE-depleted cells exhibited a positive γH2A signal, while only ∼10% uninduced cells were positive for the γH2A signal (Fig. 4B). Subsequently, we performed the ChIP experiment using the γH2A antibody and hybridized the ChIP product with telomere and tubulin probes (Fig. 4C). Significantly more γH2A was associated with the telomere chromatin after depletion of PolIE, while the same amount of γH2A was associated with the tubulin chromatin before and after depletion of PolIE (Fig. 4D). Therefore, depletion of PolIE resulted in an increased amount of DNA damage at the telomere. In addition, we analyzed the γH2A ChIP product by quantitative PCR using primers specific to the active *VSG2*, silent *VSG16* and *mVSG531*, the 70 bp repeats in the active ES (70 bp BES), and the 70 bp repeats in a silent ES (70 bp telo) (54). As a control, we also examined whether γH2A is associated with rDNA and *SNAP50*, a chromosome internal gene. As shown in Fig. 4E, depletion of PolIE resulted in an increased amount of γH2A associated with the subtelomere chromatin but not with chromosome internal loci.

**Figure 4.**
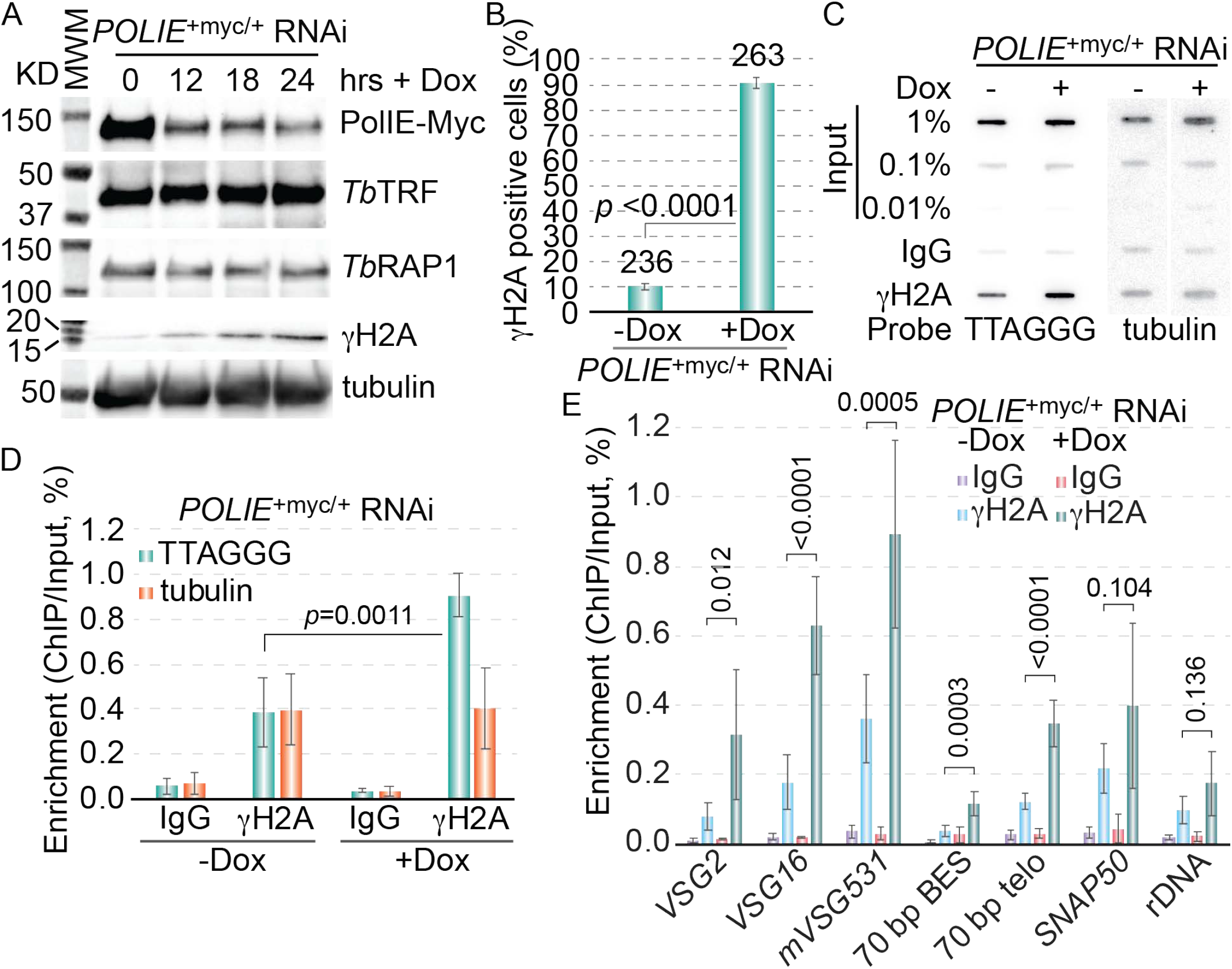
Depletion of PolIE leads to an increased amount of DNA damage. (A) Depletion of PolIE-myc in *POLIE*^+myc/+^ RNAi cells did not change *Tb*TRF or *Tb*RAP1 protein levels but increased the γH2A level. The myc Ab 9E10, a *Tb*TRF rabbit Ab (36), a *Tb*RAP1 rabbit Ab (37), a γH2A rabbit Ab (43), and the tubulin Ab TAT-1 (81) were used in western analyses. (B) Quantification of percent of *POLIE*^+myc/+^ RNAi cells that are positive for the γH2A signal in IF before and after the induction of PolIE RNAi for 24 hrs. (C) Representative Southern slot blots of γH2A ChIP products. ChIP was performed in *POLIE*^+myc/+^ RNAi cells before and 24 hrs after induction of PolIE RNAi using a γH2A rabbit Ab (43) and IgG (as a negative control). (D) Quantification of Southern blotting results of γH2A ChIP products. The average enrichment of γH2A at the telomere (ChIP/Input) was calculated from five independent experiments. (E) The γH2A ChIP products were analyzed by quantitative PCR using primers specific to the active *VSG2*, silent *VSG16* and *mVSG531*, 70 bp repeats in the active ES (70 bp BES) and a silent ES (70 bp telo), and chromosome internal *SNAP50* and rDNA. The average enrichment of γH2A at each indicated locus was calculated from five to seven independent experiments. *P* values of unpaired *t*-tests are shown in (B), (D), and (E).

DSBs at or near the active *VSG* locus have been shown to be a potent inducer for VSG switching (55, 56). Therefore, we further examined whether depletion of PolIE affected VSG switching. We introduced the PolIE RNAi construct into the HSTB261 strain that was established for VSG switching analysis (57) and named the strain as S/IEi, where S stands for switching (42). VSG switching assay was performed as we did previously (41–43). This S strain (HSTB261) has two selectable markers in the active ES: *BSD* immediately downstream of the ES promotor and *PUR-TK* between the 70 bp repeats and the active *VSG2* gene (Fig. S3) (57). Switchers will invariably stop expressing the *TK* gene due to its silencing or its loss and become resistant to Ganciclovir (GCV). Because recovering VSG switchers relies on cell proliferation, we only induced PolIE RNAi for 30 hrs followed by removal of doxycycline from the medium and selection for switchers with GCV. Removal of doxycycline after 30 hrs of induction allowed cells to recover, and these cells were still responsive to doxycycline upon repeated treatment (Fig. 5A). To monitor the PolIE protein level, we tagged one endogenous PolIE allele with 13x myc to generate the S/IEi/PolIE-myc strain. In these cells, induction of PolIE RNAi led to the depletion of PolIE-myc, and removal of doxycycline resulted in the recovery of the PolIE-myc protein level (Fig. 5B, top). Subsequently, in the S/IEi cells, we found that a transient depletion of PolIE for 30 hrs resulted in an ∼3-fold higher VSG switching rate when compared to the S/ev strain with an empty RNAi construct (Fig. 5C), indicating that PolIE suppresses VSG switching. To confirm that this phenotype is specifically due to depletion of PolIE, we introduced an ectopic allele of PolIE-myc in S/IEi cells. Adding doxycycline induced both PolIE RNAi and the expression of the ectopic PolIE-myc (Fig. 5B, bottom). The VSG switching rate in S/IEi+ecPolIE-myc cells is significantly lower than that in S/IEi cells and similar to that in S/ev cells when all cells were induced by doxycycline for 30 hrs (Fig. 5C), indicating that the ectopic PolIE-myc expression suppressed the increased VSG switching rate phenotype in PolIE-depleted cells.

**Figure 5.**
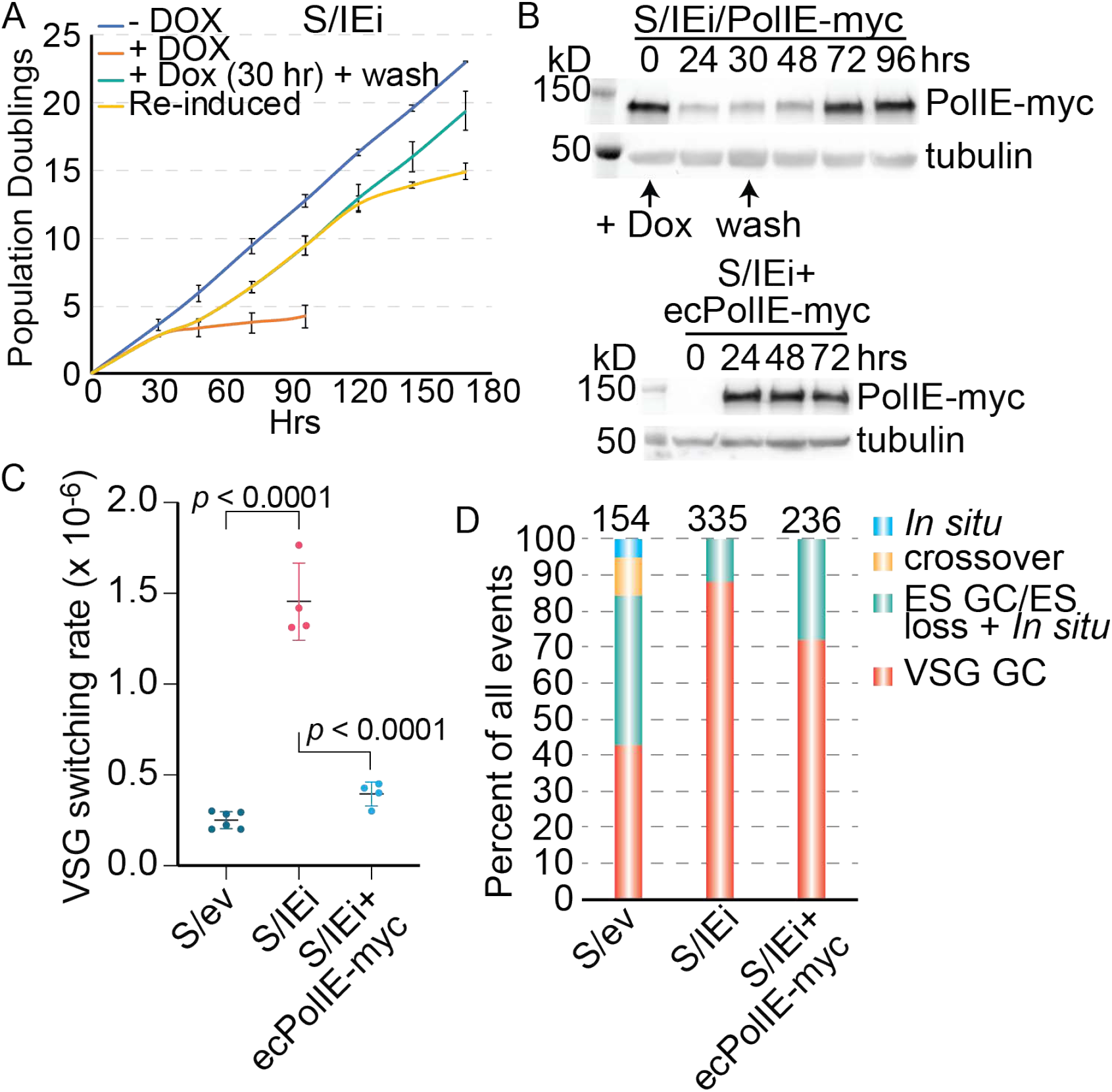
A transient depletion of PolIE leads to an increased VSG switching rate. (A) Growth curves of S/IEi cells under several conditions: without induction (-Dox), continued induction (+Dox), a transient induction (+Dox for 30 hrs followed by wash), and re-induction (after a transient 30-hr induction and wash, +Dox at 96 hrs). (B) Top, inducing PolIE RNAi for 30 hrs resulted in a transient depletion of PolIE-myc in S/IEi/PolIE-myc cells. Bottom, induced ectopic PolIE-myc expression in S/IEi+ecPolIE-myc cells. The myc Ab 9E10 and a tubulin Ab TAT-1 (81) were used in western blotting. (C) VSG switching rates in the indicated strains. *P* values of unpaired *t*-tests are shown. (D) Percent of various VSG switching mechanisms in the indicated strains. The total number of switchers characterized in each strain is listed on top of each column.

By examination of the transcription status of the *BSD* and *PUR-TK* markers in the originally active ES and by determining whether these markers and the active *VSG2* were retained in the switchers, we determined the VSG switching pathways in all obtained switchers (Fig. S3). In the S/ev control cells, a small fraction of the switchers arose from *in situ* switch (5%) and crossover (10%), while *VSG* gene conversion (43%) and ES gene conversion/ES loss + *in situ* events (42%) were more popular (Fig. 5D; Fig. S3). In contrast, in cells depleted of PolIE, *in situ* switcher and crossover events were absent, a small fraction of switchers arose from ES gene conversion/ES loss + *in situ* events (12%), while *VSG* gene conversion became the predominant switching events (88%) (Fig. 5D). Therefore, WT PolIE suppresses VSG gene conversion events by maintaining telomere integrity.

PolIE has been shown to affect VSG silencing (49). To examine PolIE’s effect on monoallelic VSG expression, we examined the transcriptomic profile in PolIE-depleted cells. Total RNA was isolated from cells where PolIE RNAi was induced for 0 or 30 hrs, and poly(A) RNA was enriched for making cDNA libraries followed by high throughput RNA sequencing analysis (Novogene Inc.). We found that the expression level of a small number of genes have been changed (either upregulated or downregulated) upon depletion of PolIE (Fig. S4A). Among the 68 upregulated genes, 44 are *VSG*s and 3 are *ESAGs* (Fig. S4C), indicating that normal VSG silencing is disrupted in cells lacking PolIE. Interestingly, three *PARP* genes encoding procyclins and transcribed by RNA Pol I were also upregulated (Fig. S4C). Procyclins are the major surface glycoproteins expressed in PF *T. brucei* that proliferates in the mid-gut of its insect vector and are silent in BF cells. Therefore, PolIE appears to affect the expression of all RNA Pol I transcribed surface antigen genes. To further validate this observation, we did quantitative RT-PCR in cells induced for PolIE RNAi for 24, 30, and 48 hrs. The active *VSG2* and rRNA expression level did not change upon depletion of PolIE, while the expression levels of all tested silent *VSGs* (including ES-linked *VSGs 3, 6, 9* and two metacyclic *VSGs*) were upregulated for nearly 10 folds after a 30-hr induction of PolIE RNAi (Fig. S4B).

To further confirm that depletion of PolIE resulted in a true VSG derepression phenotype and that different originally silent *VSGs* were derepressed simultaneously in individual cells, we performed IF. As shown in Fig. S4D, the originally silent *VSG6* and *VSG3* were both expressed after PolIE was depleted. VSG6 was more predominantly deposited on the cell surface, while VSG3 was mostly in the cytoplasm. Nevertheless, co-expression of originally silent *VSGs* strongly suggests that this is a true VSG derepression phenotype. Depletion of PolIE also resulted in downregulation of a number of genes (Fig. S4A), and more than one third of the affected genes are involved in metabolism (Fig. S4C). However, downregulated genes are much more mildly affected than the up-regulated genes (Fig. S4A).

### PolIE-depleted cells have an increased amount of T-circles and C-circles

We found that depletion of PolIE led to an increased number of VSG gene conversion events. We suspect that depletion of PolIE also affects telomere stability. *T. brucei* does not have the Non-Homologous End Joining (NHEJ) machinery (58), but HR frequently occurs at the telomere vicinity (e.g. mediating VSG switching), and intratelomeric HR can result in extra chromosomal telomere circles (T-circles, Fig. S5A, left), which frequently have nicks on both DNA strands. *T. brucei* telomere DNA also forms the T-loop structure (13), and excision of T-loop can also form T-circles (59) (Fig. S5A, left). We performed 2D gel electrophoresis, which separates circular from linear DNA molecules more clearly than 1D gel electrophoresis, followed by Southern hybridization with a telomere probe. In PolIE RNAi cells, before the induction of PolIE RNAi, there is a faint signal representing the circular DNA (Fig. 6, A, B). After depletion of PolIE, the T-circle signal is much stronger (Fig. 6, A, C), indicating that PolIE suppresses T-circle formation. We further performed the φ29 DNA polymerase-mediated telomere C-circle assay (Fig. S5, A, B) (60), which can only amplify circles with nicks/gaps on one strand. Strikingly, depletion of PolIE led to a significant increase in the amount of telomere C-circles but did not affect the telomere G-circles (Fig. 6, D, E; Fig. S5C). These observations suggest that PolIE normally suppresses telomere HR.

**Figure 6.**
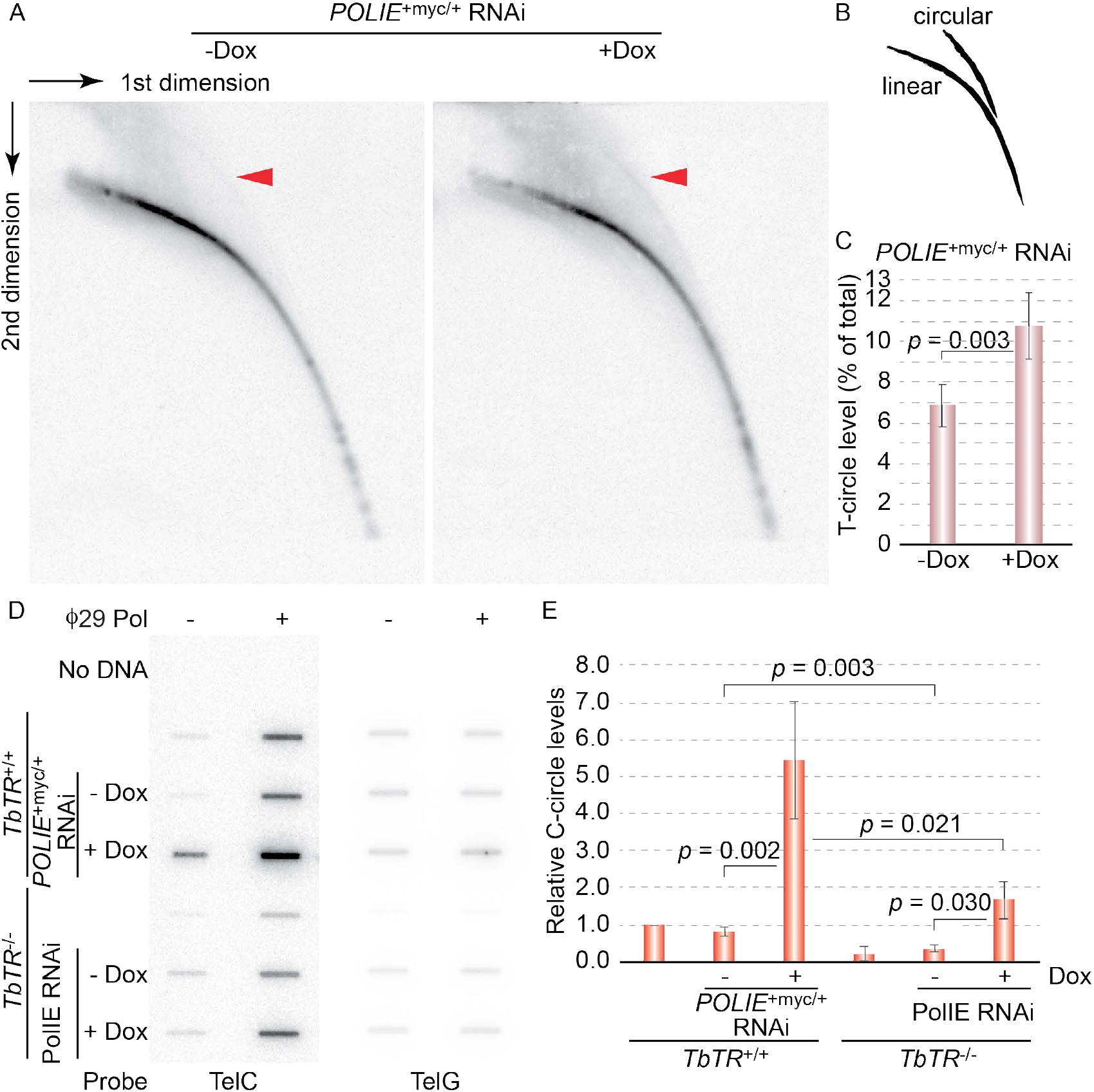
Depletion of PolIE leads to an increased amount of telomere T-circles and C-circles. (A) 2D gel electrophoresis of AluI/MboI digested genomic DNA isolated from *POLIE*^+myc/+^ RNAi cells before (-Dox) and after (+Dox) 24-hr of RNAi induction. (B) Diagram showing expected migration patterns of linear and circular DNAs in 2D electrophoresis. (C) Quantification of Southern results after 2D electrophoresis. The average T-circle amount (percent of total telomeric DNA) was calculated from six independent experiments. (D) The C-circle products from indicated cells were detected in Southern slot blotting using either a (CCCTAA)_4_ (TelC) probe or a (TTAGGG)_4_ (TelG) probe. (E) Quantification of the telomeric C-circle amount in the C-circle assay. The C-circle level in WT cells was arbitrarily set to 1, and relative C-circle levels in other cells were quantified using the WT level as a reference. The average C-circle signal level was calculated from three or four independent experiments. *P* values of unpaired *t*-tests are shown in (C) and (E).

The telomere C-circle has been identified as a hallmark of ALT cancer cells (61), which maintain the telomere length through telomere HR rather than by telomerase (62). In addition, inhibiting telomerase activity can induce cancer cells to activate the ALT telomere maintenance pathway (63). Therefore, we examined whether deleting the telomerase RNA gene (*TbTR*) could enhance the increased telomere C-circle phenotype in PolIE-depleted cells. Interestingly, there is a much lower level of telomere C-circles in *Tb*TR null cells than in WT cells (Fig. 6, D, E). This is also confirmed when comparing uninduced PolIE RNAi and *Tb*TR null/PolIE RNAi cells: deleting *Tb*TR decreased the telomere C-circle amount significantly (Fig. 6E). Therefore, deleting telomerase does not induce telomere HR directly, at least within a short time frame. In addition, depletion of PolIE in the *Tb*TR null background again induced a significant increase in the telomere C-circle level (Fig. 6, D, E), indicating that the increased amount of telomere C-circles in PolIE-depleted cells does not depend on telomerase.

### Depletion of PolIE results in an increased amount of telomere G-overhangs

The active ES-adjacent telomere is transcribed by RNA Pol I into a long non-coding telomere repeat containing RNA (TERRA) (43, 46, 64). Together with the telomere DNA, TERRA has a propensity to form the telomeric R-loop (65), a higher than WT level of which has been shown to induce more frequent telomere/subtelomere recombination in *T. brucei* (43, 46). Since PolIE depletion induced a mild VSG derepression (Fig. S4), we suspect that depletion of PolIE may also result in a higher level of TERRA. To our surprise, we detected a lower level of TERRA in PolIE RNAi cells 24 hrs after than before the induction in both northern hybridization and slot blot analyses (Fig. 7). Therefore, it is unlikely that the increased telomere/subtelomere recombination is caused by an elevated telomeric R-loop level.

**Figure 7.**
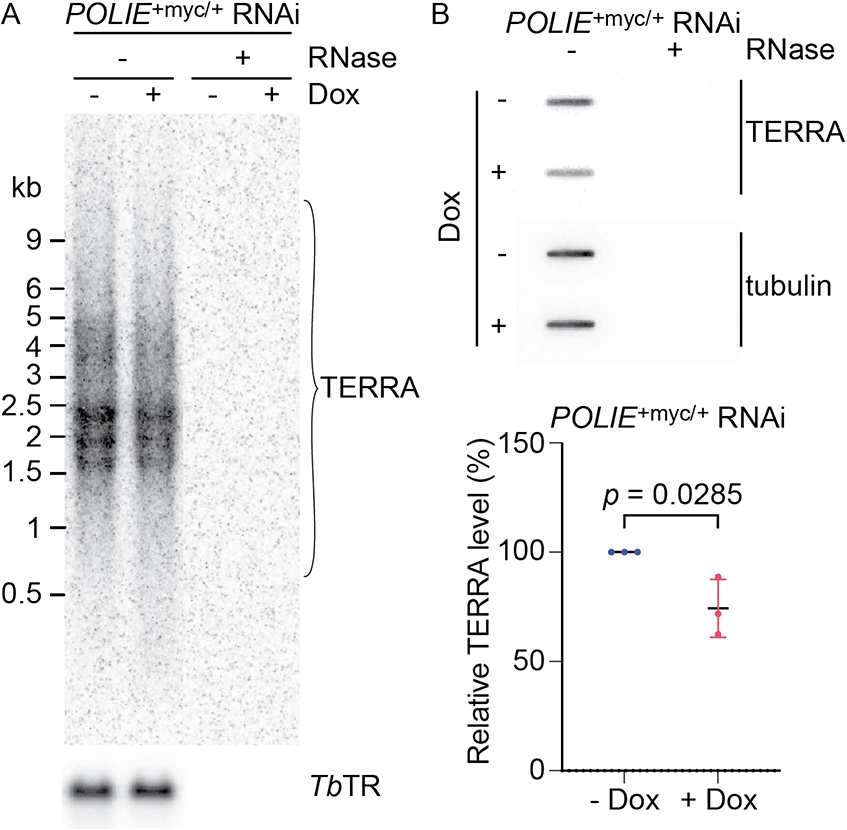
PolIE Depletion decreases the TERRA level. (A) Northern hybridization detecting TERRA in PolIE RNAi cells before (-Dox) and after (+Dox) the RNAi induction. The telomerase RNA component *Tb*TR was detected as a loading control. (B) Top, a representative northern slot blot detecting TERRA in PolIE RNAi cells. Tubulin RNA was detected as a loading control. Bottom, quantification of relative TERRA levels in PolIE RNAi cells (normalized against the *Tb*TR level). Average was calculated from three independent northern slot blot experiments. The *p* value from the unpaired *t*-test is shown.

Telomere HR can also be induced by long telomere G-overhangs, which can initiate strand invasion more efficiently. Therefore, we tested whether depletion of PolIE affected the telomere G-overhang structure in *T. brucei*. WT *T. brucei* cells have short telomere G-overhangs that are estimated to be 12-nt long or shorter (10, 11). Indeed, we only detected a very faint telomere G-overhang signal in WT cells using the native in-gel hybridization analysis (Fig. 8A). However, upon depletion of PolIE, we observed ∼20-fold more intensive telomere G-overhang signal (Fig. 8, A, C). This signal was sensitive to Exo I treatment, which degrades 3’ end single-stranded DNA specifically (Fig. 8A), indicating that the G-rich single-stranded telomere DNA is located at the end of the chromosome. In addition, only the C-rich TelC probe [(CCCTAA)_4_] detected telomere G-rich overhang signals (Fig. 8A), while the TelG probe [(TTAGGG)_4_] did not yield detectable signals (Fig. 8B), indicating that the telomere single-stranded DNA has a G-rich sequence. We also performed Pulsed-Field Gel Electrophoresis (PFGE) to separate intact *T. brucei* chromosomes and performed the same native in-gel hybridization analysis. Only G-rich telomere overhang signal was detected, and PolIE depletion greatly enhanced the telomere G-overhang signal (Fig. S6), confirming the results shown in Fig. 8. Our observations suggest that PolIE normally suppresses the telomere HR by limiting the length of telomere G-overhang. In addition, the EtBr-stained gel and the post-denaturation hybridization result showed more smeary DNA species in PolIE-depleted cells than in WT or uninduced PolIE RNAi cells (Fig. S6, left and right), further indicating that depletion of PolIE leads to an increased amount of genomic DNA degradation.

**Figure 8.**
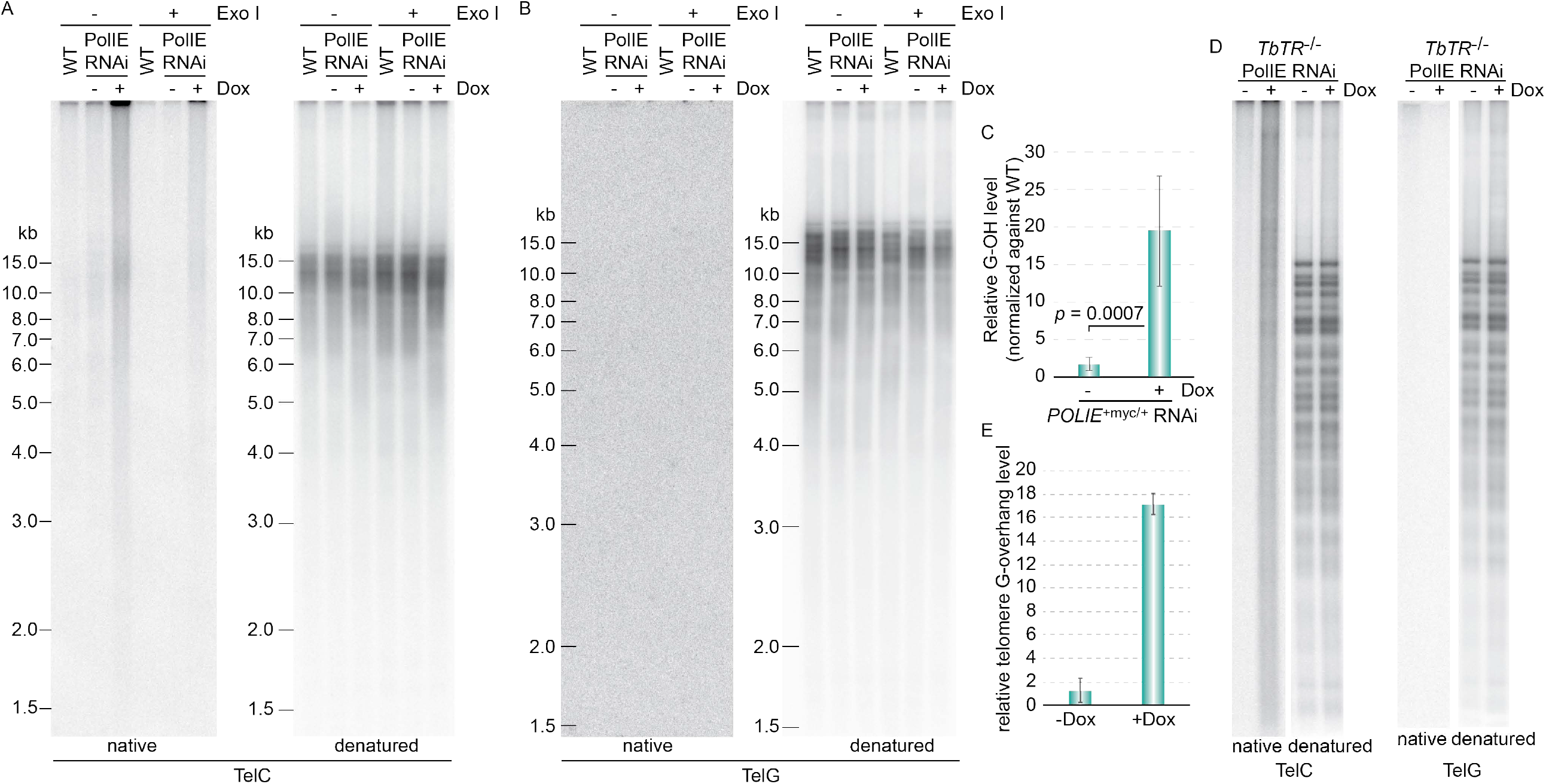
PolIE-depletion resulted in an increased level of the telomere G-overhang. Genomic DNAs were isolated from WT and *POLIE*^+myc/+^ RNAi (labeled as PolIE RNAi) cells (A & B) or from *TbTR*^-/-^ PolIE RNAi cells (D) before (-Dox) and after (+Dox) a 24-hr induction of RNAi. The genomic DNA was treated with and without ExoI (NEB), which is a 3’ single-strand DNA-specific exonuclease. In-gel hybridization was performed using a (CCCTAA)_4_ (TelC) probe or a (TTAGGG)_4_ (TelG) probe (listed below the hybridization images) first under the native condition (left) and then after denaturation and neutralization (right). (C & E) Quantification of the relative G-overhang level (using that in WT cells as a reference) in *POLIE*^+myc/+^ RNAi cells (C) and in *TbTR*^-/-^ PolIE RNAi cells (E) before (-Dox) and after (+Dox) PolIE RNAi induction. The average telomere G-overhang level was calculated from three to five independent experiments. *P* values of the unpaired *t*-test are shown in (C) and (E).

*T. brucei* telomeres have a short 3’ G-overhang, and the telomerase activity is a predominant factor for the telomere G-overhang length maintenance (10, 11). We further examined the telomere G-overhang level in *Tb*TR null/PolIE RNAi cells to determine whether the increased amount of telomere G-overhang in PolIE-depleted cells depends on the telomerase activity. Before induction of PolIE RNAi, the telomere G-overhang signal in *Tb*TR null/PolIE RNAi cells was undetectable (Fig. 8D, left). However, a significantly higher amount of the telomere G-overhang signal was observed in *Tb*TR null/PolIE RNAi cells 24 hrs after depletion of PolIE (Fig. 8D, left), indicating that the longer telomere G-overhang phenotype in PolIE-depleted cells is not dependent on the telomerase activity. In addition, quantification of the hybridization signals indicated that depletion of PolIE in the *Tb*TR null background caused ∼13-fold increase in the telomere G-overhang level, which is similar to that (∼12-fold increase) in the WT *Tb*TR background. As expected, no TelG hybridization signal was detected in *Tb*TR null/PolIE RNAi cells before and after induction.

### The subtle increase in telomere DNA synthesis in PolIE-depleted cells is telomerase-dependent

Mammalian ALT cancer cells experience frequent telomere DNA breaks that lead to elevated levels of telomere G-overhang, telomere C-circles, and telomere HR (66). We observed that PolIE depletion led to increased telomere DNA damage, telomere G-overhang length, and telomere C-circle level. Therefore, we further investigated whether PolIE is essential for telomere maintenance. Genomic DNA was isolated from PolIE RNAi cells at 0 and 24 hrs after induction and digested with MboI and AluI followed by electrophoresis on an agarose gel and subsequent Southern hybridization using a telomere repeat probe (67). Within 24 hrs of PolIE RNAi induction, no telomere length change was observed (Fig. 9A).

**Figure 9.**
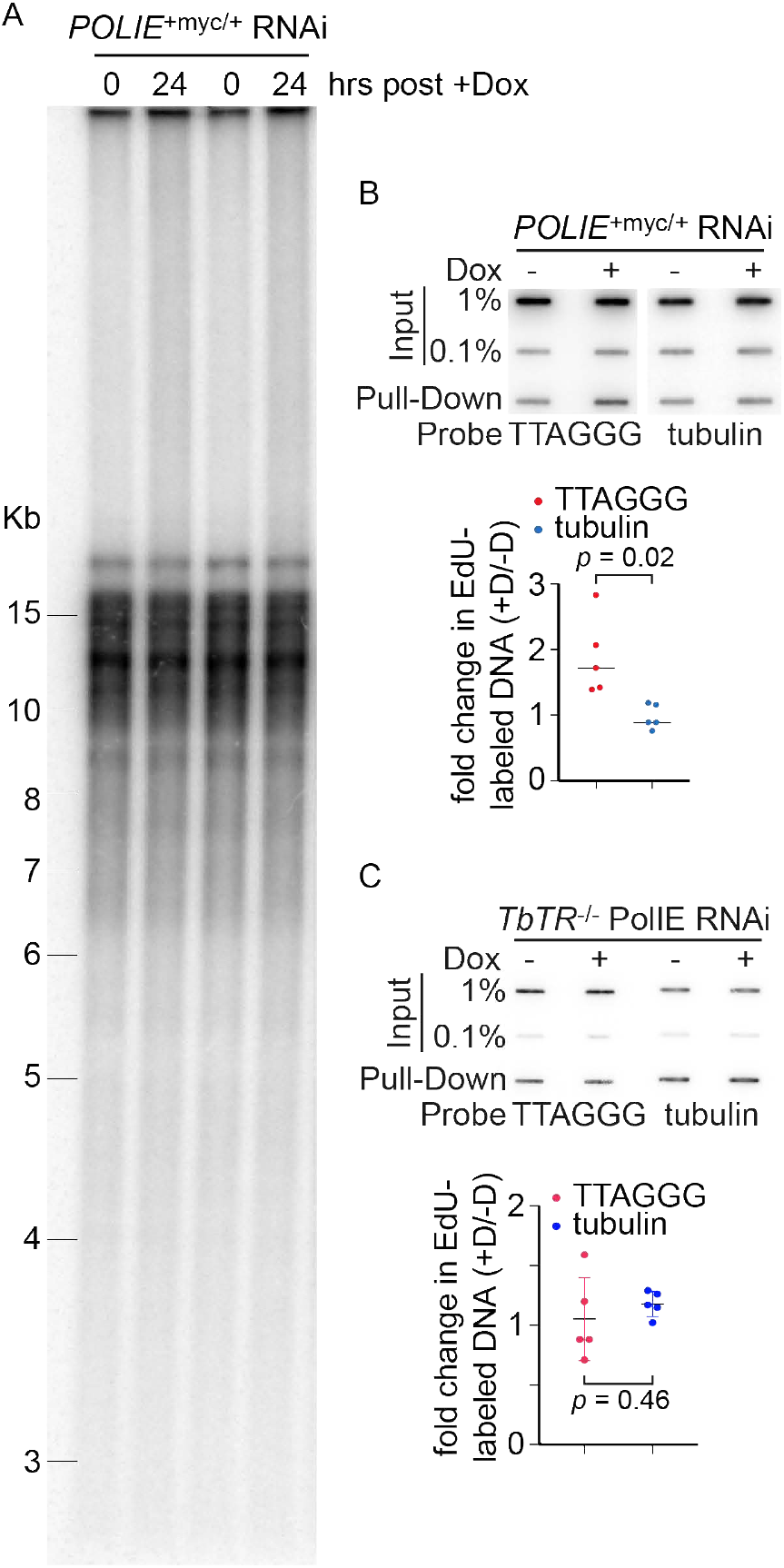
(A) Depletion of PolIE did not affect the bulk telomere length within a short time frame. Genomic DNA isolated from *POLIE*^+myc/+^ RNAi cells before (0 hr) and after (24 hr) inducing PolIE RNAi were digested with AluI and MboI and separated by agarose gel electrophoresis. Southern blotting was performed using a telomere probe. (B) and (C) In *POLIE*^+myc/+^ RNAi (B) or *TbTR*^-/-^ PolIE RNAi (C) cells, before (-Dox) and after 12-hr (+Dox) of PolIE RNAi induction, EdU-labeled nascent DNA was conjugated with desthiobiotin and pulled-down by streptavidin beads followed by Southern blotting using a telomere and a tubulin probe. The fold changes in the amount of EdU-labeled telomeric and tubulin DNA (+Dox/-Dox) were quantified and shown at the bottom in (B) and (C). The average change was calculated from five independent experiments. *P* values of unpaired *t*-tests are shown.

The telomere length changes in PolIE-depleted cells may be slow, and telomere Southern may not be sensitive enough to detect subtle changes within a short time frame. Therefore, we examined telomere DNA synthesis using an EdU labeling technique. PolIE depletion leads to cell growth arrest, which prevents proper incorporation of EdU during DNA synthesis. Hence, we performed the EdU labeling after a very brief induction of PolIE RNAi, before the cells have entered into a full growth arrest. PolIE RNAi cells were induced for 0 or 12 hrs before the cells were labeled with EdU for 3 hrs. Total genomic DNA was isolated followed by the CLICK reaction so that EdU-labeled DNA was conjugated to desthiobiotin and subsequently pulled down by streptavidin beads. The resulting EdU-labeled DNA was then hybridized with a telomere probe, and the hybridization intensities were quantified. To our surprise, telomeric DNA synthesis was at a mildly higher level in PolIE-depleted cells than in un-induced cells, while tubulin DNA replication remained the same (Fig. 9B).

Telomerase-mediated telomere synthesis is the major mechanism of telomere maintenance in *T. brucei* (33–35). To further investigate whether the subtly increased telomere DNA synthesis is dependent on telomerase, we performed the EdU labeling experiment in *Tb*TR null/PolIE RNAi cells. In the telomerase null background, depletion of PolIE no longer caused any increase in the level of telomere DNA synthesis (Fig. 9C), indicating that the subtle increase in the telomere DNA synthesis observed in PolIE-depleted cells is telomerase-dependent.

## Discussion

We identified many proteins that associate with the telomere chromatin by PICh, which is a powerful approach to isolate locus-specific chromatin (47). More than 280 proteins have been identified in both PICh and the *Tb*TRF/*Tb*TIF2 protein complex, confirming that this first application of PICh in *T. brucei* is successful and can be further applied to identify proteins that associate with other loci with repetitive sequences in *T. brucei*.

Although PolIE was originally identified as a protein that can bind to a TTAGGG repeat containing DNA oligo (48), it was not confirmed that PolIE is a telomere chromatin component. The fact that PolIE was identified in both PICh and as a component of the *Tb*TRF/*Tb*TIF2 protein complex strongly suggests that PolIE is a telomere protein. Our PolIE ChIP result further verifies that PolIE indeed associates with the telomere chromatin. In addition, we observe that PolIE co-localizes with *Tb*TRF, a good telomere marker (36), further indicating that PolIE is a telomere protein. Interestingly, although PolIE belongs to the DNA Polymerase I family in *T. brucei* (49), it has similar DNA crosslink repair functions as mammalian A-family DNA polymerase Polθ and POLN. Similar to PolIE, human DNA polymerase ν (POLN) has only the C-terminal DNA polymerase domain (68), and POLN helps repair DNA crosslinks (69, 70). In addition, Polθ, having both the C-terminal DNA polymerase domain and the N-terminal helicase domain, is also important for repairing UV-induced DNA damage in skin (71) and *POLQ* (encoding Polθ) null mouse is hypersensitive to ICL (72, 73). The function and domain structure similarities between PolIE and mammalian POLN suggest that PolIE is more homologous to POLN. Nevertheless, PolIE-depleted cells are also mildly more sensitive to EMS and MMS treatments (Fig. S2E) (49), indicating that PolIE is important for maintaining genome integrity.

We found that PolIE plays a critical role in maintaining telomere stability. Several observations indicate that PolIE suppresses HR at the telomere vicinity. First, more telomere T-circles can be detected in PolIE-depleted cells than in WT cells. Second, PolIE-depletion leads to an increased amount of telomere C-circles. Third, a transient depletion of PolIE results in many more VSG gene conversion-mediated VSG switching events. Therefore, PolIE clearly helps maintain telomere stability by suppressing telomere HR. Mammalian Polθ has also been shown to promote alternative NHEJ (MMEJ) at dysfunctional telomeres and translocations at non-telomeric genome loci but suppresses HR-mediated DNA damage repair (52). On the other hand, mammalian POLN does not seem to have any telomere-specific functions. In this sense, PolIE appears to have functions more similar to the mammalian Polθ.

Most strikingly, we discover that PolIE plays a critical role in maintaining a proper telomere 3’ G-rich overhang structure. Depletion of PolIE greatly increases the telomere 3’ G-rich overhang length, suggesting that PolIE regulates factor(s) contributing to the telomere G-overhang structure, including the telomerase-mediated G-strand synthesis, the C-strand fill-in, and the 5’ telomere end resection by exonuclease(s) (27). We previously showed that WT *T. brucei* cells have very short telomere G-overhangs (∼ 12 nts) (10). In addition, in telomerase null *T. brucei* cells, the telomere 3’ G-overhang length is significantly diminished (11). Interestingly, depletion of PolIE leads to a higher level of the telomerase-mediated telomere synthesis, indicating that PolIE normally suppresses the action of telomerase at the telomere. The telomerase-mediated telomere G-strand extension no doubt contributes to the longer telomere G-overhangs observed in PolIE-depleted cells. However, this does not seem to be the only contributing factor. In the *Tb*TR null background, the telomere G-overhangs are much longer after PolIE depletion, indicating that this phenotype is not completely telomerase-dependent. However, whether PolIE affects telomere C-strand fill-in and/or 5’ telomere resection will need further investigation.

Interestingly, PolIE appears to maintain telomere stability using different mechanisms than *Tb*RAP1 and *Tb*TRF. Both *Tb*RAP1 and *Tb*TRF suppress the TERRA and the telomeric R-loop levels, which in turn helps maintain telomere integrity and stability (43, 46). However, we find that depletion of PolIE results in a lower level of TERRA. It is possible that the long telomere G-overhangs in PolIE-depleted cells form a G-quadruplex structure, which blocks transcription elongation of RNA Pol I and results in a lower level of TERRA transcription. However, more experiments will be necessary to test this hypothesis.

*T. brucei* does not have any NHEJ mechanism (58), but telomere/subtelomere HR is a realistic threat to telomere stability, as HR-mediated VSG switching is frequently observed as a major VSG switching mechanism (41–44, 55–57, 74–76). Therefore, it is significant to discover that *T. brucei* has multiple mechanisms to suppress telomere HR, mediated by various telomere proteins including *Tb*RAP1 (43), *Tb*TRF (46), and PolIE.

## Materials and Methods

### *T. brucei* strains

Procyclic form (PF, at the insect stage) *T. brucei* WT cells (Lister 427) were transfected with pSK-*Tb*TRF-ko-*Hyg* (36) and pSK-*Tb*TIF2-ko-*BSD* (42) to delete one allele of *TbTRF* and *TbTIF2*, respectively. The resulting cells were transfected with SacII digested pSK-F2H-*Tb*TRF-*Pur*-tar and pSK-*Tb*TIF2-F2H-Phleo-tar (42) to insert the FLAG-HA-HA (F2H) tag to the N-terminus of the remaining *TbTRF* allele and to the C-terminus of the remaining *TbTIF2*, respectively. The resulting *TbTRF*^F2H+/-^*TbTIF2*^+F2H/-^cells were used for the 2-step IP of the *Tb*TRF/*Tb*TIF2 protein complex.

All bloodstream form (BF) *T. brucei* strains used in this study were derived from VSG2-expressing Lister 427 cells that express the T7 polymerase and the Tet repressor (Single Marker, aka SM) (77). The HSTB261 strain is derived from SM and specifically designed for VSG switching assay (57), which we renamed as the S strain for easier reference (42). SM cells, *Tb*TIF2^+F2H/+^(42), and the S cells were transfected with pSK-PolIE-myc13-Hyg-tar to tag one endogenous allele of *PolIE* with a C-terminal tag including 13 repeats of myc. The SM cells expressing PolIE-Myc from one of its endogenous loci (*PolIE*^+myc/+^) were transfected with pZJMβ-*Tb*TRF (36) and pZJMβ-PolIE to generate inducible *Tb*TRF RNAi and PolIE RNAi strains, respectively. The S and S/PolIE-myc cells were transfected with pZJMβ-PolIE to generate S/IEi and S/IEi/PolIE-myc strains. S/IEi cells were transfected with pLew100v5-PolIE-myc to generate the S/IEi + ecPolIE-myc strain. *TbTR*^-/-^cells (34) were transfected with pZJMβ-PolIE to generate the *TbTR*^-/-^PolIE RNAi strain. SM and *PolIE*^+myc/+^cells were transfected with pSK-PolIE-ko-BSD to generate *PolIE*^+/-^ and *PolIE*^+myc/-^ cells.

### Plasmids

A 500 bp genomic DNA fragment upstream of the *TbTRF* gene, the *Puromycin resistance* gene (*PUR*), the α/β tubulin intergenic sequence, the F2H-tag, and a 500 bp fragment of the *TbTRF* gene (encoding the N-terminus) are inserted into pBluescript SK in this order to generate pSK-F2H-*Tb*TRF-*Pur*-tar.

A 400 bp *PolIE* ORF fragment (at the C-terminus), a myc_13_ tag, the α/β tubulin intergenic sequence, the *Hygromycin resistance* gene (*HYG*), and a 500 bp genomic DNA fragment downstream of the *PolIE* gene were inserted into pBluescript SK in this order to generate pSK-PolIE-myc13-Hyg-tar.

A 470 bp DNA fragment at the N-terminus of *PolIE* ORF and a 520 bp DNA fragment at the C-terminus of *PolIE* ORF were inserted into pZJMβ (78) in tandem to generate pZJMβ-PolIE RNAi construct.

The *Blasticidin-resistance* gene (*BSD*) flanked by genomic DNA fragments upstream and downstream of the *PolIE* gene, respectively were inserted into pBluescript SK to make pSK-PolIE-ko-BSD.

### Nuclear Fractionation and 2-Step Immunoprecipitation

5 × 10^10^ *TbTRF*^F2H+/-^ *TbTIF2*^+F2H/-^ PF cells were harvested by centrifugation followed by snap freezing in liquid nitrogen. After thawing cells, the cell pellet was resuspended in the hypotonic buffer (10 mM HEPES pH 7.9; 10 mM KCl; 2.5 mM MgCl_2_; 1 mM EDTA; 1 mM DTT; Complete protease (Roche), 1 mM PMSF, 1 µg/ml Leupeptin, 0.5 mg/ml TLCK, and 1 µg/ml Pesptatin A) and incubated on ice for 10 minutes, followed by adding NP-40 to a final concentration of 0.2%. Cells were homogenized in a glass douncer until ∼80% of the cells were broken. The lysate was overlaid on top of 0.3 mL of sucrose buffer (0.8 M sucrose in hypotonic buffer) and centrifuged at 7 krpm for 10 minutes at 4 ºC. After removing the top cytosolic and sucrose layers, the nuclear fraction was resuspended in the TBS buffer (50 mM Tris•HCl, pH7.4; 420 mM NaCl; Complete protease (Roche), 1 mM PMSF, 1 µg/ml Leupeptin, 0.5 mg/ml TLCK, and 1 µg/ml Pesptatin A) and digested with Benzonase on ice for 20 minutes followed by centrifugation at 13 krpm for 10 minutes at 4 ºC. The cell lysate was then diluted with 50 mM Tris-HCl, pH 7.4 so that the final concentration of NaCl reaches 150 mM. The cell lysate was pre-cleared with Dynabeads Protein G (ThermoFisher) then incubated with Anti-FLAG M2 magnetic beads (Sigma) for 3 hrs at 4 ºC. The IP product was washed 3 times with TBS-T (50 mM Tris-HCl, pH7.4; 150 mM NaCl, 0.1% Tween 20) and eluted at room temperature with 0.6 mg of the FLAG peptide. The eluate was incubated with Dynabeads Protein G conjugated with the HA monoclonal antibody 12CA5 (MSKCC Monoclonal Ab Core) for 2 hrs at 4 ºC. The IP product was washed 3 times with TBS-T, eluted with 0.1 M glycine, pH 2.2 at 56 ºC followed by neutralization with 1 M Tris•HCl pH 9.5. The final eluate was concentrated with StrataClean resin (Agilent) before extracted using 2X SDS buffer and separated in a 10% SDS-PAGE gel (Bio-RAD). The gel was stained with colloidal Coomassie Brilliant Blue followed by mass spectrometry analysis (Cleveland Clinic Lerner Research Institute Proteomics and Metabolomics Core).

### Proteomics of Isolated Chromatin Segments (PICh)

PICh was performed according to (47) and (79) with modifications. 5 × 10^10^ WT PF cells were harvested and washed with 1 x PBS and HLB (10 mM HEPES pH 7.9; 10 mM KCl; 2.5 mM MgCl_2_; 1 mM EDTA; 1 mM DTT; Complete protease (Roche), 1 mM PMSF, 1 µg/ml Leupeptin, 0.5 mg/ml TLCK, and 1 µg/ml Pesptatin A). Cells were resuspended in HLB, incubated on ice for 10 min, and then homogenized with 5 strokes using a glass douncer. After adding 15% paraformaldehyde in HLB, 15-20 more strokes were performed. The lysate was centrifuged at 15,000 g in a swing bucket centrifuge at room temperature for 20 min, and the pellet was resuspended in 3% paraformaldehyde/1xPBS with rotation for 15 min followed by 3 washes using 1 x PBS. The pellet was then washed with the sucrose buffer (0.3 M Sucrose; 10 mM HEPES-NaOH, pH 7.9; 1% Triton-X100; 2 mM MgOAc), resuspended in the sucrose buffer and homogenized in a douncer with 20 strokes, then centrifuged at 15,000 g for 20 min at room temperature. Subsequently, the pellet was washed with 1 x PBS/0.5% Triton X-100, resuspended in 1 x PBS/0.5% Triton X-100/200 µg/mL RNAse A with rotation for 2 hrs. After two washes with 1 x PBS and one wash with HSLB (10 mM HEPES-NaOH, pH 7.9; 100 mM NaCl; 2 mM EDTA; 1 mM EGTA; 0.2% SDS; 0.1% Sodium Sarkosyl; 1 mM PMSF), the pellet was resuspended in HSLB and incubated at room temperature for 15 mins, followed by sonication for 13 cycles (30 sec on/off) at maximum intensity in a Bioruptor 300 (Diagenode). Samples were heated to 58 ºC for 5 mins in a Thermomixer, then spun down at 20 krpm for 15 minutes at room temperature. The supernatant was rotated for 2 hrs with pre-equilibrated High Capacity Streptavidin Agarose Resin slurry (Thermo). Then the mixture was centrifuged for 2 mins and the supernatant was desalted by filtration with a Sephacryl S-400 HR column (GE Healthcare), centrifuged at 16 krpm, followed by adding SDS to a final concentration of 0.02%. The resulting desalted soluble chromatin was divided into 2 equal aliquots to which either a telomere specific [TelG, 5’ TTAGGGTTAGGGTTAGGGTTAGGGT 3’] or scramble (Scr, 5’ GATGTGGATGTGGATGTGGATGTGG 3’) LNA probe were added. Chromatin and LNA probes were hybridized in a thermocycler (25 ºC/3 min, 71 ºC/9 min, 38 ºC/60 min, 60 ºC/3 min, 38 ºC/30 min, 25 ºC final temperature), then centrifuged, immunoprecipitated, eluted, and TCA concentrated as published (79). Crosslink reversal was done by suspending the pellet in LDS NuPAGE sample buffer + 0.5 M 2-Mercaptoethanol and incubating at 99 ºC for 25 mins. Protein samples were resolved in 4-12% NuPAGE Bis-Tris gels (ThermoFisher), stained with colloidal Coomassie Brilliant Blue and stored in sealed packages until mass spectrometry analysis.

### EMS Clonogenic Survival Assay

BF PolIE RNAi cells were grown with or without doxycycline for 24 hours then incubated with 2 mM of ethylmethanesulfonate (EMS; Sigma) or DMSO (vehicle) for two hours. Cells were washed three times with warm media and plated at a concentration of 1 cell/well in 96-well plates followed by incubation at 37 ºC. Wells were checked for growth daily under the microscope until the 12^th^ day after plating. Survival was calculated as a percentage of EMS treated wells presenting growth in relation to the respective DMSO controls.

### UV and Cisplatin sensitivity tests

*PolIE*^+myc/+^ PolIE RNAi cell were first incubated with and without doxycycline for 12 hrs then diluted to a concentration of 1.5 × 10^6^cells/ml. Cells were irradiated by 0, 50 J m^-2^, or 100 J m^-2^UV using the Stratalinker® UV Crosslinker (Stratagene) or incubated with 0, 10, and 20 µM cisplatin (Sigma) for 1 hr. Cells were washed free of doxycycline and cisplatin after the treatment and cell growth was monitored daily with necessary dilutions. Cell relative growth was calculated using the following formula for cells incubated with and without doxycycline: population doubling (UV/cisplatin treated)/population doubling (untreated). The average was calculated from three independent experiments.

### 2-dimensional gel electrophoresis to detect telomeric T-circles

Genomic DNA was isolated from PolIE RNAi cells incubated with and without doxycycline for 24 hrs the same way as published previously (67). 5 µg of MboI/AluI digested genomic DNA were separated by 2-dimensional gel electrophoresis according to (80). Specifically, DNAs were separated in the first dimension in a 0.4% agarose 1 x TBE gel without ethidium bromide under 40 volts for 18 hours at room temperature. After the 1^st^ dimension electrophoresis, the gel was incubated in 1 x TBE/0.3 µg/ml ethidium bromide for 20 minutes followed by washing with 1 x TBE. DNA migration was verified by a long-wave UV light. DNAs were then excised from the gel, transferred to a second 1.1% agarose gel with 1 x TBE/0.3 µg/ml of ethidium bromide. DNAs were electrophoresed under 150 volts at 4°C for 5 hrs. Subsequently, DNA was transferred to a Nylon membrane (GE Healthcare) by blotting followed by hybridization with a telomere probe.

### The C-circle (G-circle) Assay

The φ29-mediated rolling-circle assay was performed according to (60) with minor modifications. 8 µg of *T. brucei* genomic DNA was first digested by AluI and MboI (40 U each) and 100 ng of DNase-free RNase A, then by λ Exonuclease and Exonuclease I to remove dsDNA. 20 ng of the resulting DNA was incubated with 7.5 U φ29 DNA polymerase (NEB) in reaction buffer [1 µg/µl BSA, 0.05% Tween 20, 0.5 mM dATP, 0.5 mM dGTP (or 0.5 mM dCTP for detecting G-circles) and 0.5 mM dTTP, 1 x φ29 Buffer] at 30 °C for 8 h, then heat-inactivate the enzyme at 65 °C for 20 min. The reaction products were dot-blotted and UV cross-linked onto a Hybond N nylon membrane (GE Healthcare) followed by hybridized at 50 °C with end-labeled (CCCTAA)_4_ to detect C-circles or (TTAGGG)_4_ to detect G-circles.

### Native in-gel hybridization to detect the telomere G-overhang structure

Genomic DNA was digested with MboI and AluI followed by the treatment with or without Exo I. Equal amount of ExoI treated and non-treated DNA were separated by agarose gel electrophoresis. The agarose gel, with separated DNA fragments inside, was dried at room temperature without denaturation. The Gel was then hybridized with an end-labeled (CCCATT)_4_ or (TTAGGG)_4_ probe at 50°C. After extensive wash and exposing the hybridized gel to a phosphorimager, the gel was denatured, neutralized, and hybridized with the same oligo probe at 55°C followed by exposure to a phosphorimager. The hybridization signals were quantified using ImageQuant. The telomere G-overhang level was calculated by dividing the amount of native hybridization signal by the amount of post-denaturation hybridization signal.

Alternatively, DNA plugs were prepared according to (36). Intact *T. brucei* chromosomes were separated by pulse-field gel electrophoresis according to (42). The agarose gel, with separated chromosome DNA, was dried and hybridized with the (CCCATT)_4_ or (TTAGGG)_4_ probe the same way as described above. The hybridization signals were calculated the same way as described above.

### Telomere Southern to estimate telomere length

was performed the same way as described in (67).

### EdU-labeling

Exponentially growing BF *T. brucei* cells (0.7-0.9 × 10^6^ cells/ml) were incubated with 150 µM 5-ethynyl-2’-deoxyuridine (EdU) (Click Chemistry Tools) for 3 hrs before genomic DNA was isolated. DNA was sonicated to 400-1000 bp fragments. EdU-labeled DNA fragments were conjugated with desthiobiotin using Click chemistry regent (2 mM desthiobiotin-Azide, 100 mM/500 mM CuS04/THPTA, 50 mM Na-Ascorbate, 100 mM HEPES pH 7, and 10% DMSO). The desthiobiotin conjugated DNA was pulled-down using streptavidin beads (ThermoFisher) and eluted from the beads using biotin. The eluted DNA was dot-blotted and UV cross-linked onto a Hybond N nylon membrane (GE Healthcare) and hybridized with a telomere (or a tubulin) probe at 65°C. The blot was exposed to a phosphorimager and the signals were quantified using ImageQuant.

## Acknowledgements

We thank Dr. Keith Gull for the TAT-1 antibody. We thank the Li lab members for their comments on the manuscript.

