## Supplemental materials for "*Trypanosoma brucei* PolIE suppresses telomere recombination"

**Supplemental figure legends**

Figure S1. Schematic diagrams of two approaches to identify telomere protein components. Left, nuclear extract was prepared from PF *TbTRF*$^{F2H+/-}$ *TbTIF2*$^{+F2H/-}$ cells. The FLAG monoclonal antibody M2 (Sigma) was used in the first IP. The FLAG IP product was then IPed with the HA monoclonal antibody 12CA5 (MSKCC Antibody & Bioresource Core Facility). The final IP product was separated by PAGE and the whole lane was analyzed by mass spectrometry (Cleveland Clinic Lerner Research Institute Proteomics and Metabolomics Core). WT PF cells were treated exactly the same way and the sequential IP product was also analyzed by mass spectrometry as a negative control. Right, WT PF cells were crosslinked with formaldehyde before the cell lysate was sonicated and incubated with the $(TTAGGG)_4$-containing LNA probe conjugated with the desthiobiotin. The LNA-hybridized telomere chromatin was pulled down by streptavidin beads and eluted by Biotin. A control LNA containing a random sequence was used as a negative control. The final pull-down products were separated by PAGE and analyzed by mass spectrometry.

Figure S2. (A) Growth curves of *POLIE*$^{+myc/+}$, *POLIE*$^{+myc/-}$, *POLIE*$^{+/+}$, and *POLIE*$^{+/-}$. Average population doublings were calculated from three independent experiments. (B) and (C) ChIP experiments were performed in *POLIE*$^{+myc/+}$ (B) and *POLIE*$^{+myc/+}$ *Tb*TRF RNAi (C) cells using the myc monoclonal antibody 9E10 (MSKCC Antibody & Bioresource Core Facility) and a *Tb*TRF rabbit antibody (36). IgG was used as a negative control. The ChIP product was analyzed by slot blot hybridization using a telomere and a tubulin probe. (D) Quantification of ChIP slotblot hybridization results using the tubulin probe in *POLIE*$^{+myc/+}$ *Tb*TRF RNAi cells. (E) PolIE-depleted cells are more sensitive to EMS treatment. Survival of EMS-treated cells was calculated as percent of untreated cells before and after induction of PolIE RNAi by doxycycline. Average was calculated from three independent experiments. *P* value (unpaired *t*-test) is shown.

Figure. S3. A schematic diagram of the *in vitro* VSG switching assay (57) [adapted from (43)]. The S strain (parent, top left (57)) has a *blasticidin resistance (BSD)* marker immediately downstream of the active ES promoter (long arrow) and a *puromycin resistance (PUR)-thymidine kinase (TK)* fusion gene inserted between the 70 bp repeats and the active *VSG2* gene. Other ESs are silent. In an *in situ* switch (bottom left), the active ES becomes silenced while a previous silent ES becomes expressed. In crossover/telomere exchange events, the active *VSG2* gene and its neighboring DNA sequences change place with a silent *VSG X* (frequently also in a ES). In *VSG* gene conversion events, a silent *VSG X* is duplicated into the active ES to replace the active *VSG2* gene, which is lost. In ES gene conversion events, a whole silent ES is duplicated to replace the originally active ES, which is lost. In ES loss coupled with an *in situ* switch (ES Loss + *in situ*), the originally active ES is lost and a different ES is expressed. In each scenario, the expected resistance (R) or sensitive (S) phenotypes to 100 μg/ml blasticidin, 2 μg/ml puromycin, 5 μg/ml GCV, and the *VSG2* and *BSD* genotypes (tested by PCR analyses) are listed. +, gene is present; -, gene is absent.

Figure S4. Depletion of PolIE affects VSG monoallelic expression. (A) Volcano plot shows differential gene expression in PolIE RNAi cells before and after induction. (B) Quantitative RT-PCR results in PolIE RNAi cells show derepression of several BF ES-linked *VSGs* and metacyclic ES-linked *mVSGs*. (C) Different classes of genes are upregulated or downregulated upon depletion of PolIE. (D) IF analysis shows that depletion of PolIE results in co-expression of two originally silent VSGs in individual cells, confirming a true VSG derepression phenotype.

Figure S5. (A) Schematic diagram showing that intratelomeric excisions can leads to T-circles (left). Diagram of telomeric C and G-circles are shown on the right. (B) Schematic diagram showing the principle of the C-circle amplification assay. (C) Depletion of PolIE leads to an increased amount of telomere C-circles. The C-circle amplification products were analyzed by slotblot hybridization using a C-rich telomere probe. Average fold difference in C-circle amount (+Dox/-Dox) is calculated from three independent experiments.

Figure S6. Native in-gel hybridization analysis to examine the telomere G-overhang structure. DNA plugs were prepared from PolIE RNAi cells before (-) and after (+) the induction. Undigested chromosomes were separated by PFGE. After electrophoresis, the gel was dried at room temperature followed by hybridization with end-labeled TelC or TelG probes under the native condition. After exposing the gel to a phosphorimager, the gel was denatured, neutralized and again hybridized with the same TelC or TelG probes. The Ethidium bromide-stained gel is shown on the left. The native in-gel hybridization result is shown in the middle. The hybridization result after the denaturation is shown on the right.

Tonini et al. Figure S1

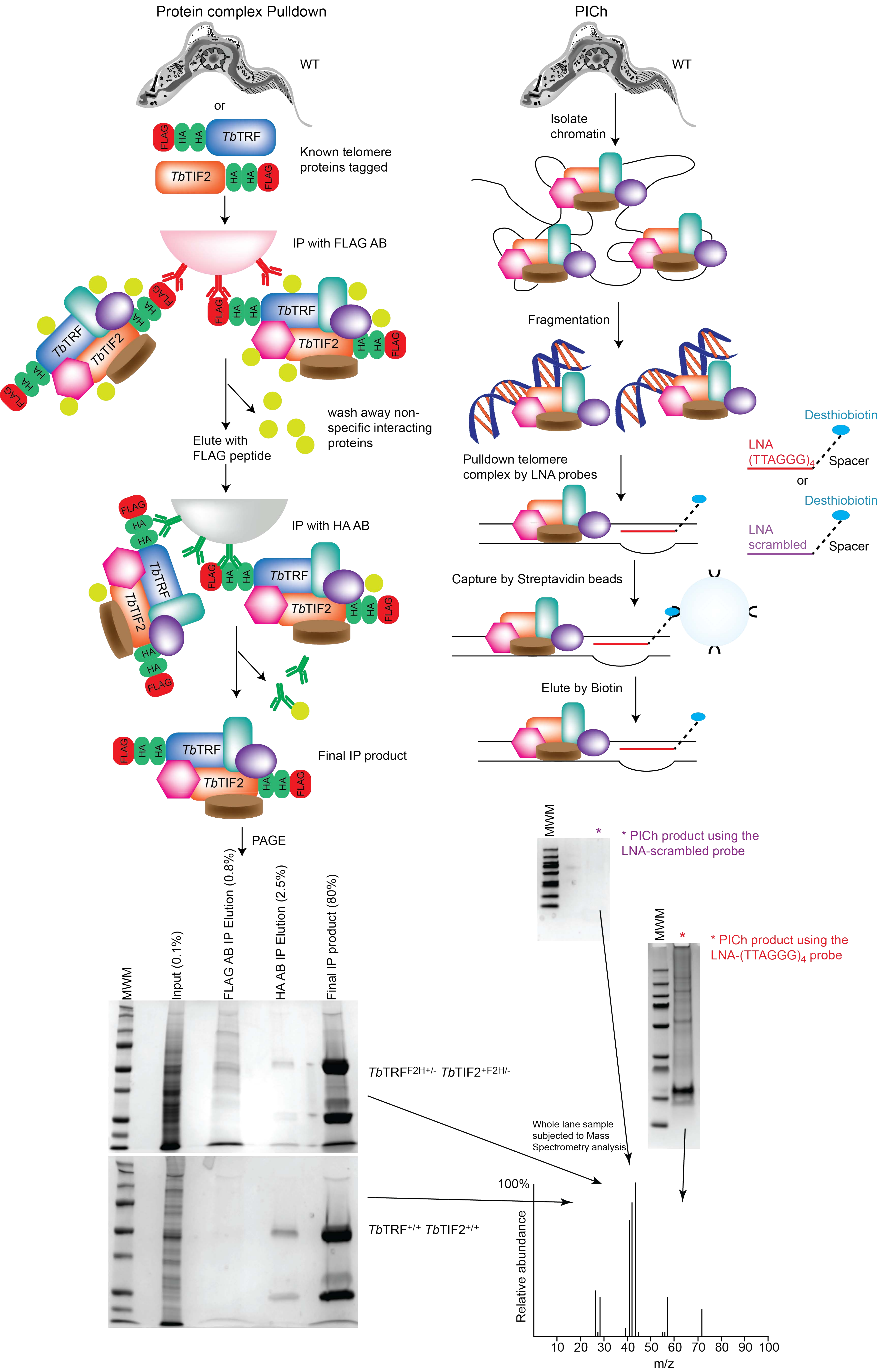

Tonini et al. Figure S2

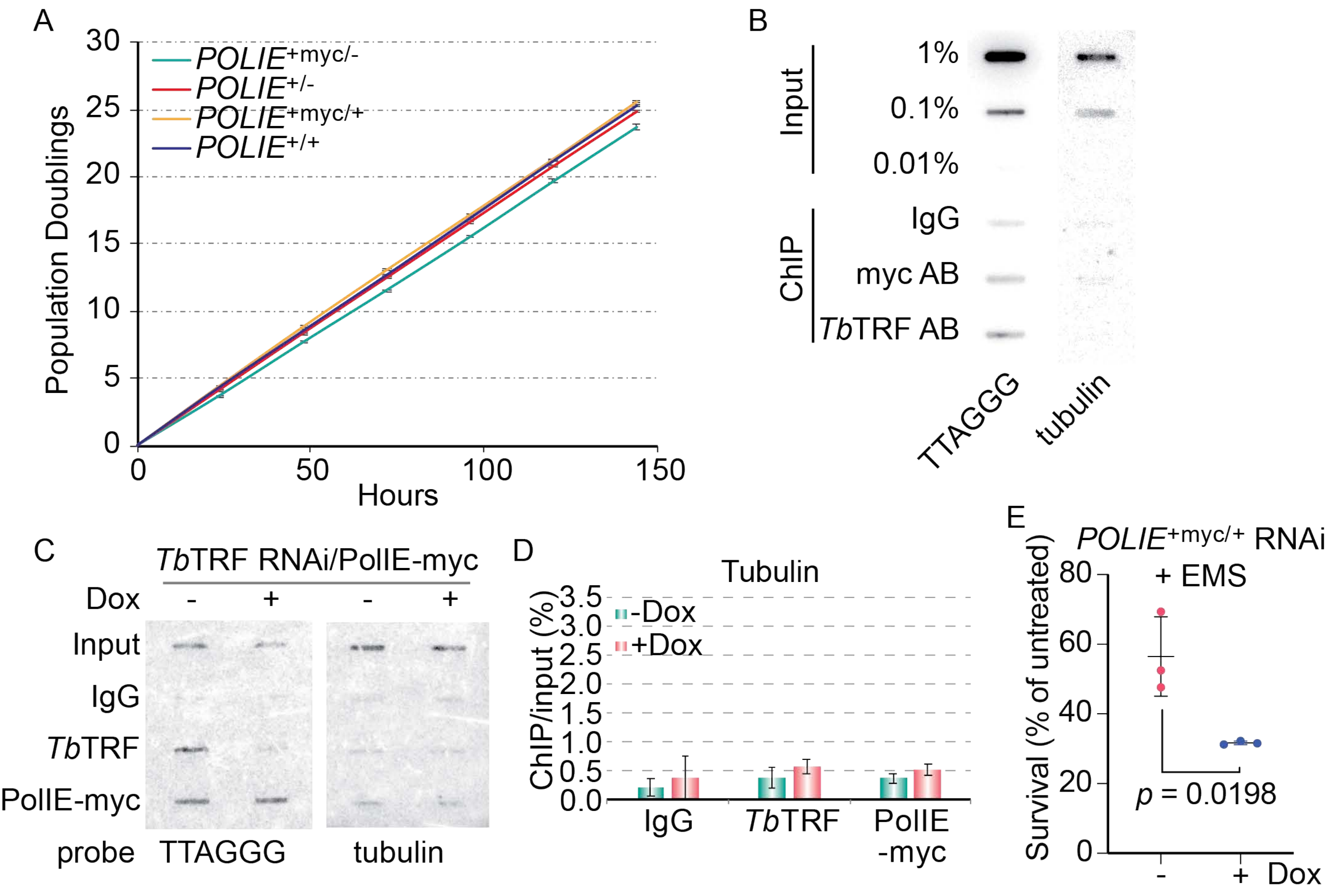

Tonini et al. Figure S3

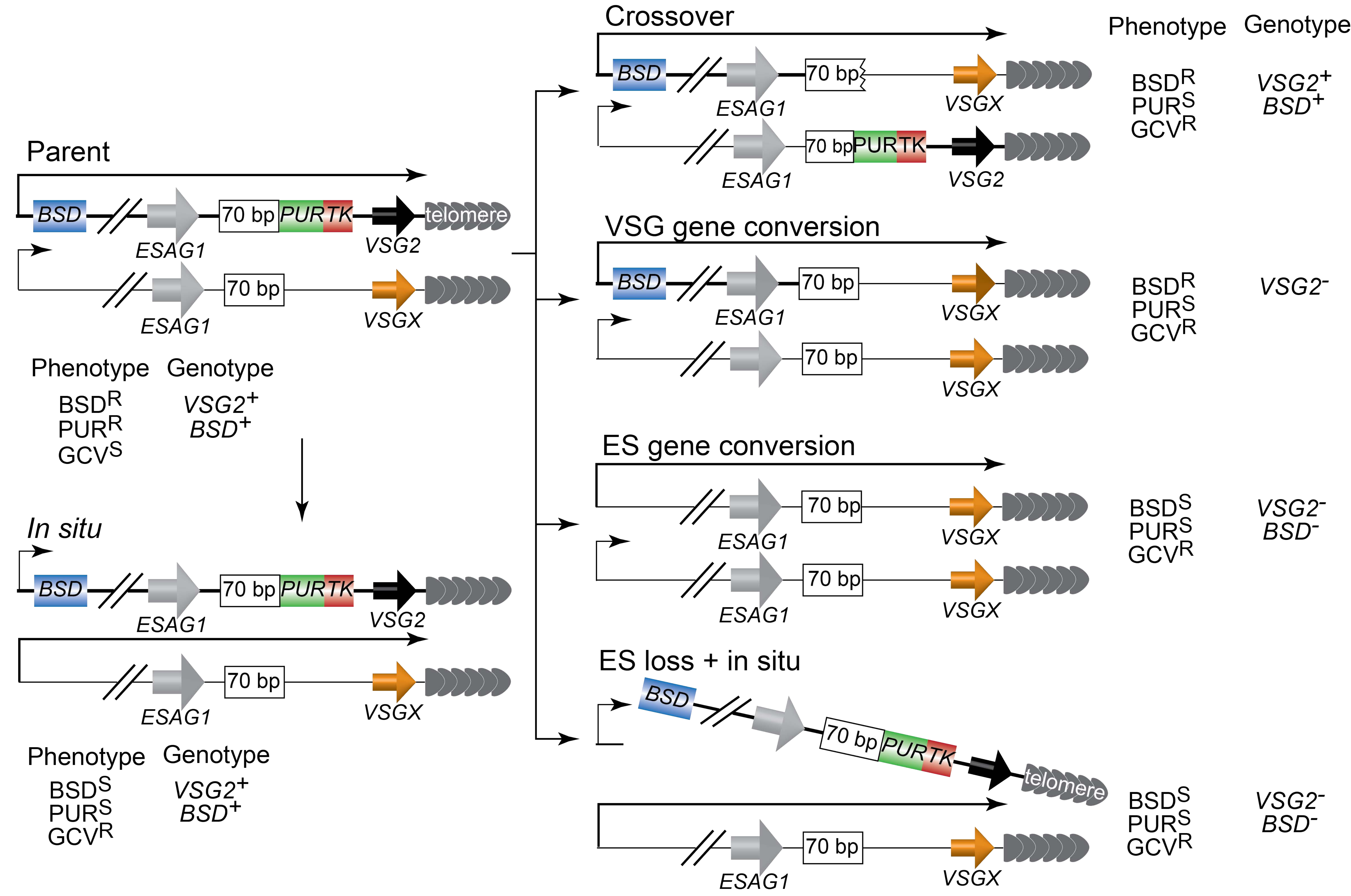

Tonini et al. Figure S4

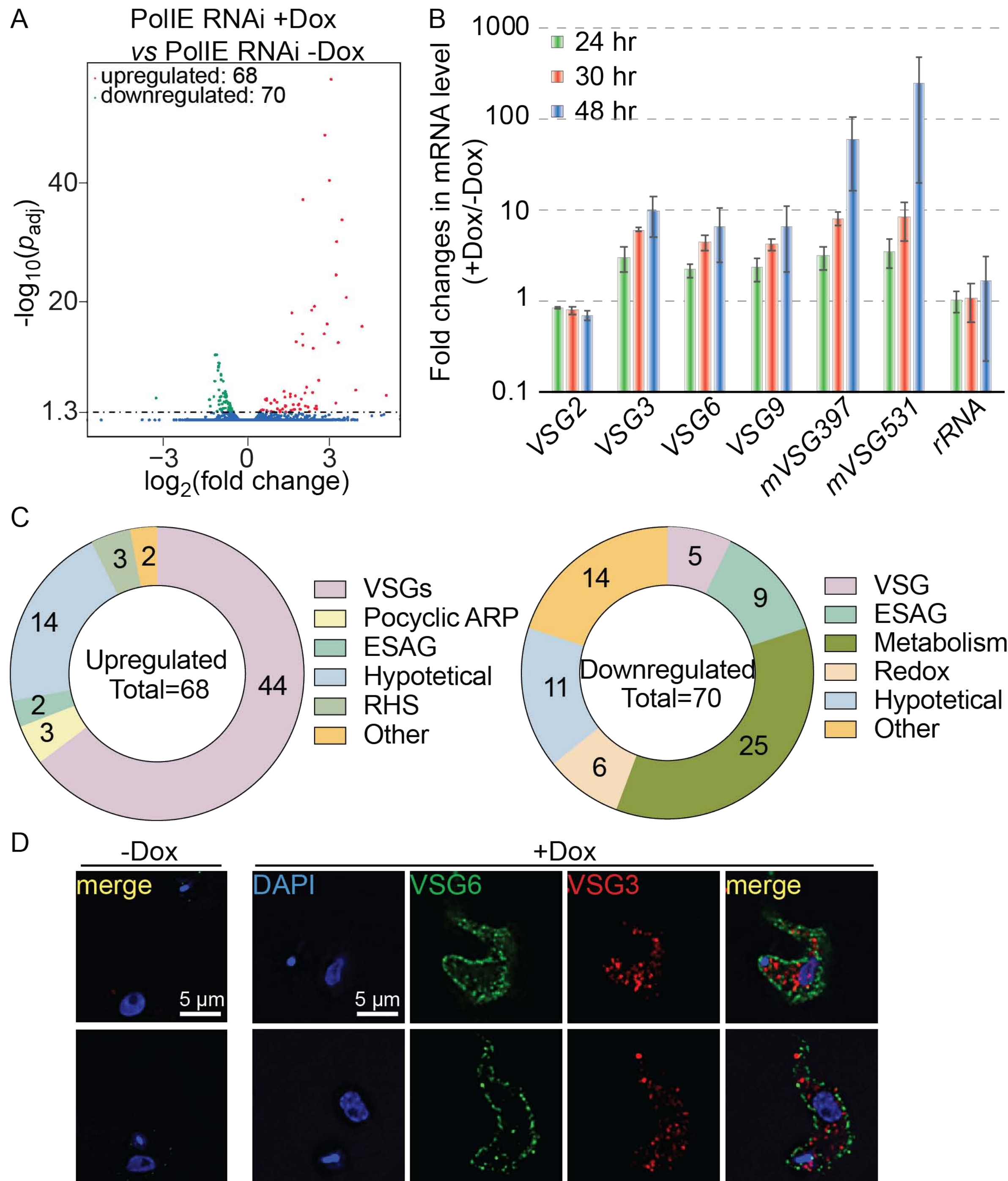

Tonini et al. Figure S5

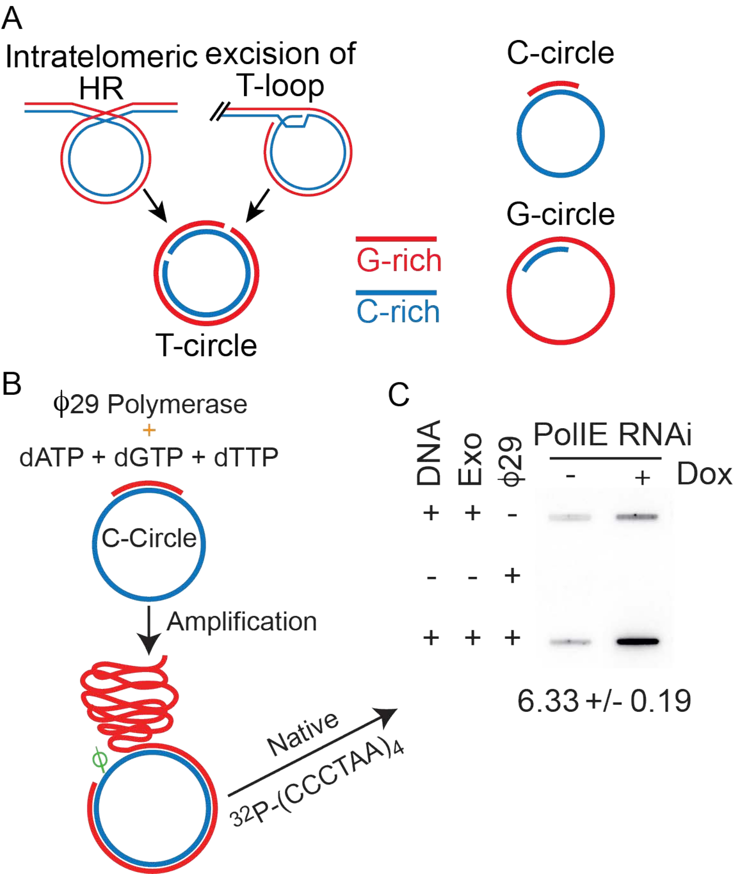

Tonini et al. Figure S6

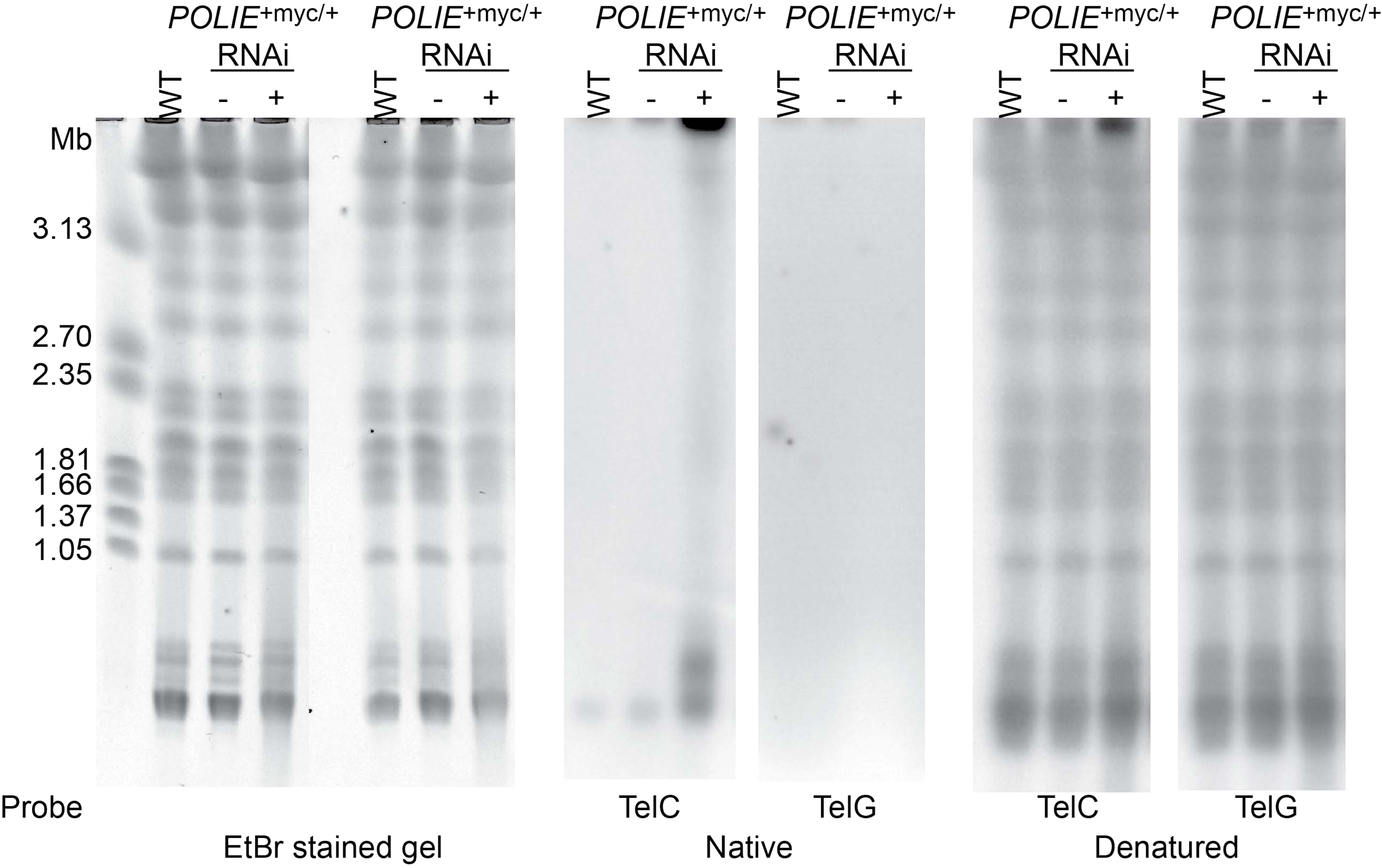
